# Manipulation of the nucleoscaffold potentiates cellular reprogramming kinetics

**DOI:** 10.1101/2023.03.12.532246

**Authors:** Benjamin A. Yang, André Monteiro da Rocha, Isabel Newton, Anna Shcherbina, Sing-Wan Wong, Paula M. Fraczek, Jacqueline A. Larouche, Harrison L. Hiraki, Brendon M. Baker, Jae-Won Shin, Shuichi Takayama, M. D. Thouless, Carlos A. Aguilar

## Abstract

Somatic cell fate is an outcome set by the activities of specific transcription factors and the chromatin landscape and is maintained by gene silencing of alternate cell fates through physical interactions with the nuclear scaffold. Here, we evaluate the role of the nuclear scaffold as a guardian of cell fate in human fibroblasts by comparing the effects of transient loss (knockdown) and mutation (progeria) of functional Lamin A/C, a core component of the nuclear scaffold. We observed that Lamin A/C deficiency or mutation disrupts nuclear morphology, heterochromatin levels, and increases access to DNA in lamina-associated domains. Changes in Lamin A/C were also found to impact the mechanical properties of the nucleus when measured by a microfluidic cellular squeezing device. We also show that transient loss of Lamin A/C accelerates the kinetics of cellular reprogramming to pluripotency through opening of previously silenced heterochromatin domains while genetic mutation of Lamin A/C into progerin induces a senescent phenotype that inhibits the induction of reprogramming genes. Our results highlight the physical role of the nuclear scaffold in safeguarding cellular fate.

## INTRODUCTION

All mammalian cells strictly regulate gene expression to retain their identity and function through genomic organization of chromatin^1,2^. Recent studies have shown that the spatial segregation of transcriptionally active and inactive chromatin domains is achieved in part through physical interactions with subnuclear structures at the nuclear periphery^3–6^. The periphery of the nucleus is bounded by the inner and outer nuclear membrane and the nucleoskeleton, a complex mesh of type V intermediate filaments called lamins^7^. Nuclear lamins are comprised of A-type (*LMNA*) and B-type (*LMNB1, LMNB2*) isoforms^8^, and their expression is lineage-restricted such that all somatic cells express at least one B-type lamin^7^ while A-type lamins are lacking from embryonic stem cells and restricted to differentiated cells^9^. A-type lamins also scale in abundance according to cell type^10–12^ and tissue stiffness^13^. Accordingly, the animate functions of the genome and cell-type specificity^10,14^ are primarily mediated through dynamic interactions with lamins, whereby genes associated with alternate lineages are condensed into transcriptionally inactive heterochromatin and physically tethered to the nuclear periphery in lamina-associated domains (LADs)^12,15–17^. LADs are 0.1-10 Mb in size^17^, cover more than one-third of the human and mouse genomes^18^, and are associated with transcriptionally inactive domains^15^. During stem cell differentiation or activation, hundreds of repressed genes in LADs detach from lamins to increase in expression and promote cell fate transitions^10,14,19,20^. Thus, nuclear lamins are a safeguard for cellular fate and a gateway to previously repressed genes.

The loss of functional lamins in laminopathies such as Hutchinson-Gilford Progeria Syndrome (HGPS), which is driven by mutations in the *LMNA* gene^21^, results in disruption to LADs, abnormal nuclear morphologies, loss of peripheral heterochromatin, and premature aging^22,23^. HGPS and other laminopathies have also displayed altered nuclear stiffness^24^ and impairments in the ability to resist mechanical loads, producing susceptibility to changes in chromatin conformation^25^. The ability of the nucleus to resist mechanical deformation is specified in part by separate contributions from lamins and heterochromatin domains, in which chromatin is responsible for the elastic response at small deformations (<3μm), while A-type and B-type lamins regulate the elastic and strain-stiffening responses, respectively, in response to larger mechanical stimuli^26–28^. Taken together, these results suggest that lamins sit at the nexus between chromatin organization, nuclear stiffness, and intracellular mechanical signaling, but how these proteins influence cellular fate remains underexplored.

In this article, we evaluate how changes in the composition of nuclear lamina impact the stability of cellular fate using somatic cell reprogramming as a model system, whereby a differentiated cell fate is erased and a new cellular identity is adopted. We compare the reprogramming kinetics^29^ of healthy fibroblasts, HGPS fibroblasts possessing permanently dysfunctional *LMNA*, and fibroblasts that transiently downregulate Lamin A/C to evaluate how chromatin state, elasticity of the nucleus^16^, and cell fate plasticity are coupled. We first show how loss of functional Lamin A/C proteins affects morphology, and chromatin accessibility. Next, we assess differences in the mechanical properties of a nucleus using a novel micro-mechanical cell-squeezing device. Finally, we demonstrate that aberrant nuclear morphology and associated changes in the mechanical elasticity associate with changes in reprogramming efficacy and kinetics. To understand the molecular mechanisms that confer these changes, we mapped transcriptional dynamics of single cells across conditions, and show that Lamin A/C knockdown produces varied TF-binding dynamics. These results shed light on how mechanical properties of a nucleus focalized through lamins contribute to chromatin organization and cellular fate.

## RESULTS

### Disruptions in Nucleo-Scaffold Affect Nuclear Morphology and Heterochromatin

To evaluate the effect of disruptions to the nuclear lamina on nuclear morphology, we examined 3 different systems. First, we utilized dermal fibroblasts possessing a heterozygous point mutation in exon 11 of the *LMNA* gene (c. 1824C>T; p.Gly608Gly) obtained from patients with HGPS^22^. This mutation activates a cryptic splice donor site, resulting in the synthesis of a truncated, immature form of Lamin A/C called progerin^22^ (**Supp. Figure 1A,B**) that has been shown to produce aberrant nuclear morphology^30^ and loss of peripheral heterochromatin^23^. We confirmed the presence of progerin in HPGS fibroblasts (**Supp. Figure 1A)** and compared HGPS fibroblasts to healthy human dermal fibroblasts (Controls) as well as fibroblasts that had temporarily reduced levels of Lamin A/C (*LMNA* KD). Knockdown of Lamin A/C was achieved through delivery of Dicer-substrate small interfering RNAs^31^ (DsiRNAs) targeting *LMNA* in lipid-nanoparticles. Following *LMNA* KD, we observed >75% reduction of mRNA levels and significant reductions in *LMNA* levels (**Figure 1A,B; Supp. Figure 1C,E,F**) with minimal impact on cell proliferation (**Supp. Figure 1D-F**). Immunostaining for Lamin A/C and two different markers associated with constitutive heterochromatin (H3K9me3 and HP1a) in HGPS, *LMNA* KD, and control fibroblasts revealed that both HGPS and *LMNA* KD fibroblasts displayed alterations in nuclear morphology when compared to unmodified controls (**Figure 1A**). Specifically, multiple protrusions or cavities were observed along the contour of the nucleus for HGPS and *LMNA* KD cells when compared to controls, likely reflecting punctate areas of localized protein depletion (**Figure 1C,D**). In addition, we observed ruptures of the nuclear membrane in Lamin A/C staining for *LMNA* KD (**Figure 1A,C**), indicating elevated instability in nuclear architecture. *LMNA* KD also resulted in increased H3K9me3, independent of the heterochromatin protein 1a (HP1a) (**Figure 1A,B**), which was in contrast to HGPS fibroblasts that displayed reductions in H3K9me3 and HP1a^5^. These results are congruent with previous studies^5,32^ and the notion that modifying the amount and type of lamins engenders alterations to nuclear morphology and heterochromatin.

**Figure 1.**
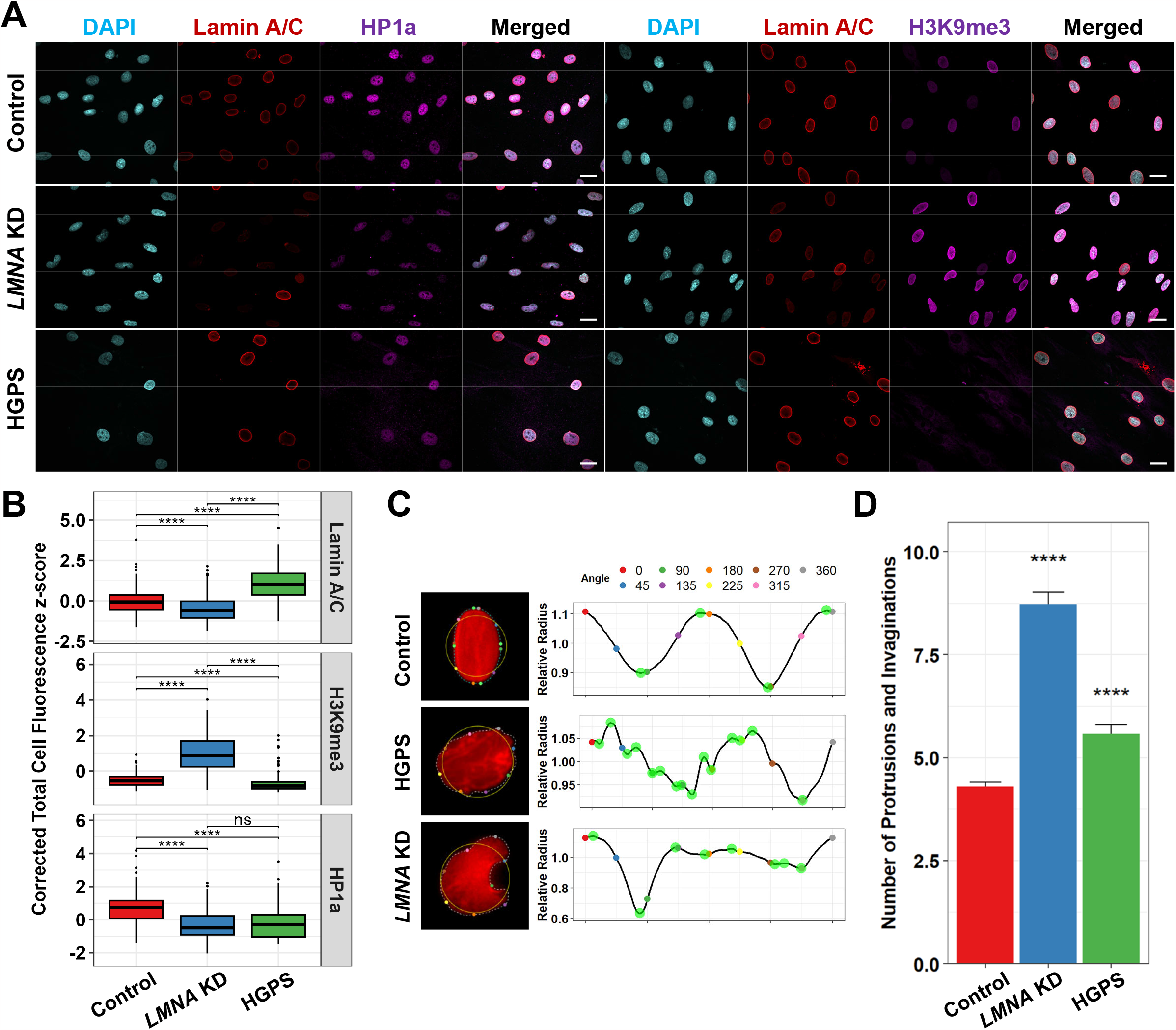
Modifications to the Nucleo-Scaffold Engender Alterations in Nuclear Morphology. Immunofluorescence analysis of Lamin A/C, H3K9me3, and HP1a in DAPI-counterstained nuclei from control, *LMNA* knockdown, Hutchinson-Gilford Progeria Syndrome (HGPS), and reprogrammed (hiPSC) human dermal fibroblasts. **A)** Representative images and **B)** mean fluorescence intensities per nucleus (z-scores) are shown. Scale bars are 25 μm. Statistical comparisons are Student’s *t*-tests with Holm multiple comparison correction. **C)** Representative images of fibroblast nuclei stained for Lamin A/C and corresponding nuclear perimeter curves for quantitation of nuclear invaginations and protrusions. **D)** Average number of nuclear protrusions and invaginations per condition. Statistical comparisons are two-sided Mann-Whitney U-tests. Data are shown as mean +/- SEM. ****p<0.0001.

### Nuclear Scaffold Alterations Modulate Chromatin Accessibility and Transcription Factor Binding

Inducing somatic cells to reprogram or change fate requires the concerted re-expression and binding of previously silenced transcription factors (TFs), which physically untether from the nuclear envelope back towards interior nuclear compartments^33^. Previous research has shown that HGPS cells display reductions in heterochromatin and increased chromatin accessibility in LADs^34^, but less is known about how transient loss of lamins impacts chromatin accessibility and gene accessibility within LADs. To probe further whether transient knockdown of *LMNA* facilitates heterochromatin detachment that results in changes in transcription factor binding (**Figure 2A**), we utilized an assay for transposase-accessible chromatin followed by high-throughput sequencing^35,36^ (omni-ATAC-Seq) on control fibroblasts, *LMNA* KD and human embryonic stem cells (hESCs). We generated 11,779,630 read pairs on average for each sample, resulting in 150,008 enriched sites across all conditions. Principal component analysis (PCA) of enriched sites (**Figure 2B**) revealed distinct patterns between each condition in which *LMNA* KD and control fibroblasts clustered separately from hESCs along principal component 1 and hESCs lie in between control and *LMNA* KD along principal component 2. Partitioning each dataset according to their shared and unique sites (**Figure 2C**) revealed that unique sites were mostly distal to gene promoters (**Figure 2D**). To identify patterns in altered chromatin accessibility, we performed gene set enrichment analysis of unique sites between control and *LMNA* KD fibroblast using the GREAT toolbox^37^. This analysis revealed that *LMNA* KD promoted increased accessibility around genes associated with a variety of pro-reprogramming factors, including heparan sulfate synthesis, which is highly upregulated in ESCs and is critical to their self-renewal^38^, G-protein coupled glutamate receptor signaling, which promotes the undifferentiated state in cultured ESCs^39,40^, Activin receptor signaling, which regulates stem cell self-renewal^41^, and PIWI-associated RNA (piRNA) biogenesis, which are uniquely expressed in reprogrammed pluripotent cells^42^. PIWI-associated RNAs have also been shown to contribute to H3K9me3 deposition at transposable elements^43^, in support of elevated H3K9me3 expression after Lamin A/C knockdown. We next identified differences in TF binding using differential TF footprinting analysis^44^. This analysis revealed enrichments of pro-reprogramming motifs in *LMNA* KD that are implicated in the epithelial-to-mesenchymal transition (EMT) during embryogenesis^45^ (*SNAI1/2/3*) in addition to Notch signaling (*HES1*), *YY2*, which regulates post-transcriptional expression of reprogramming factors (*OCT4, KLF4, SOX2*, and *c-MYC* – OKSM)^46^, and *HINFP*, which regulates cell proliferation and repression of transposable elements^47^. In contrast, control fibroblasts were enriched for AP-1 family members, which are known to guard the somatic genome from reprogramming^48,49^, and *TEAD4*, which is known to regulate transcription networks supportive of reprogramming^50^ (**Figure 2F)**. These results suggest transient knockdown of the nuclear scaffold creates a chromatin landscape that results in accessibility of previously silenced transposable elements^43,51,52^ and pathways and TF-binding associated with stem cell self-renewal (**Figure 2E**).

**Figure 2.**
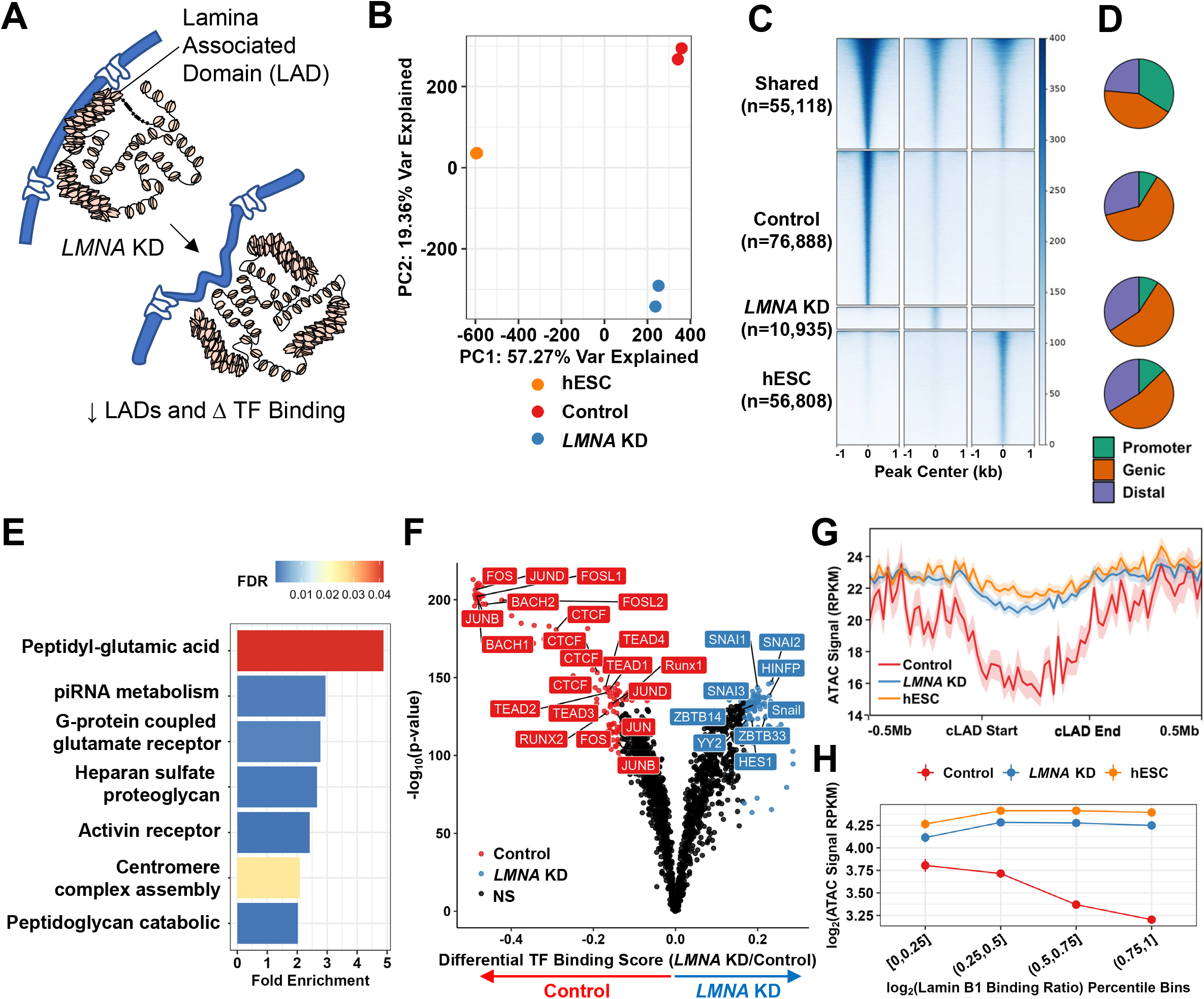
Changes in Chromatin Accessibility Patterns Following Nucleo-Scaffold Manipulation Permit Alternative Transcription Factor Binding. **A)** Schematic showing how knockdown of Lamin A/C results in changes in lamina-associated domains (LADs) and opening of chromatin for transcription factor (TF) binding necessary for reprogramming to pluripotency. **B)** Principal component analysis of enriched chromatin accessibility sites identified by omni-ATAC-Seq. **C)** Heatmaps of chromatin accessibility in shared and unique sites in a 2 kb region centered on the nearest TSS. **D)** Annotations of enriched sites relative to promoter regions. **E)** Enriched GO terms identified through GREAT analysis in accessible sites unique to *LMNA* knockdown fibroblasts compared to control fibroblasts. **F)** Differential TF footprinting between control and *LMNA* knockdown fibroblasts. **G)** Chromatin accessibility signal (in RPKM) around constitutive lamina-associated domains (cLADs). **H)** Chromatin accessibility in log_2_(RPKM) within and around cLADs as a function of Lamin B1 binding percentile bins.

To further evaluate whether the increases in chromatin accessibility of transposable elements were derived from LADs, we integrated datasets of constitutive lamina-associated domains (cLADs), which are conserved across cell types and species^53^. We found that while control fibroblasts showed decreased ATAC signal within cLADs, hESC and *LMNA* KD fibroblasts both showed opening of cLADs (**Figure 2G,H**). Integrating these results shows loss of lamins promotes alterations in chromatin organization and increased access to previously silenced motifs required for adoption of a new cellular fate.

### Development of Micro-Mechanical Squeezing Assay to Assess Nuclear Deformations

Nuclear mechanical properties are typically assessed in bulk by confining cells between parallel plates^54^, stretching an elastic substrate on which cells are adhered^55,56^, or using atomic force microscopy (AFM)^57–59^, micropipette aspiration^60^, micropost arrays^61^, or microfluidic devices^62^. While these approaches have begun to tease apart the relationship between nuclear morphology and stiffness in three-dimensions, they typically rely on highly specialized equipment and setups. We improved upon an easily configurable micro-engineered PDMS-based device^64^ with custom stretching mechanisms to precisely trap and reproducibly deform single cells with detailed three-dimensional optical read outs enabled by confocal microscopy (**Figure 3A**). The microfluidic device uses a substrate of poly(dimethylsiloxane) (PDMS) that contains micro-patterned notches rigidly attached to a thin film of hard PDMS (h-PDMS) (**Supp. Figure 2A**). Application of tensile strain to the device generated micro-cracks^63^ (**Supp. Figure 2B**) owing to both a lower toughness and three-fold increase in the modulus of the thin hPDMS layer compared to the underlying substrate (**Supp. Figure 2C**). The resultant cracks can then be opened and closed in a reproducible manner by varying strain of the device^64^ according to the thickness of the h-PDMS layer (**Supp. Figure 2D,E**). Three-dimensional confocal imaging of the channels when the device was strained and relaxed showed the fully reversible nature of the channel cross-section (**Figure 3B**).

**Figure 3.**
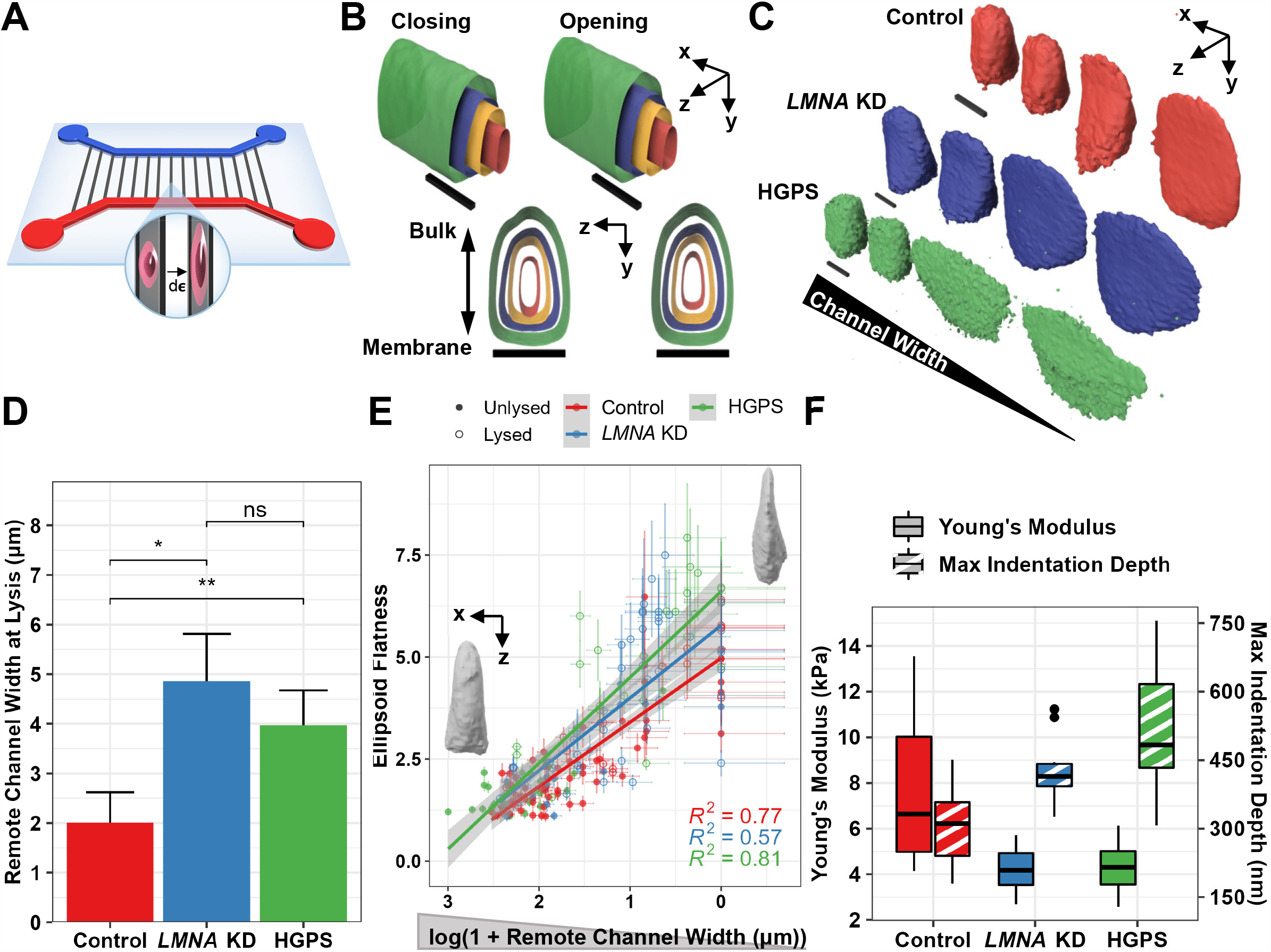
Loss of Functional Lamins Reduces Nuclear Resistance to Lysis and Compression. **A)** Schematic of device whereby single cells are mechanically trapped and gradually deformed through release of device strain (dε). **B)** Confocal reconstructions of microchannel cross-sections as device strain is released (closing) and re-applied (opening). Cross-sections are colored green, blue, yellow, and red, in order of decreasing device strain and channel width. Scale bars (black bars) are 1 μm × 1 μm × 10 μm. **C)** Confocal reconstructions of nuclei ordered by decreasing channel width (left to right) and colored by cell type. Scale bars (black bars) are 1 μm × 1 μm × 10 μm. **D)** Average channel width distal to cells at the point of nuclear lysis. **E)** Flatness of ellipsoids fitted to each nucleus as a function of channel width distal to the cell. Lysis status is marked by open and closed circles. Cell type-specific trends are fitted with linear regression models. (HGPS, n=14; Control, n=20; *LMNA* KD, n=16; hiPSC, n=17). Representative confocal reconstructions at lower and upper ellipsoid flatness values are shown. **F)** Young’s moduli and maximum probe indentation depths of nuclei across conditions as measured through atomic force microscopy of intact cells above the nucleus. (Control, n=15; HGPS, n=11; *LMNA* KD, n=10). Statistical comparisons are unpaired Mann-Whitney U-tests. **p<0.01, ***p<0.001.

To test the system’s ability to resolve morphological changes during compression, hydrogel spheres with comparable sizes (∼25 μm diameter) and mechanical properties (2.4 kPa) to cells were fabricated from alginate polymers and conjugated to rhodamine using a cross-junction microfluidic device^65^ (**Supp. Figure 3A,B**). The hydrogel spheres were loaded into the system and their deformation observed at varying strains using confocal microscopy (**Supp. Figure 3C**). Bead deformation was assessed by fitting an ellipsoid at each channel width. The alginate hydrogel was observed to deform linearly^65^ in proportion to decreases in channel width and partially recover its undeformed morphology after the channel was reopened (**Supp. Figure 3C-E**). The direction of bead deformation was primarily parallel to decreasing channel width with minimal changes in morphology along the length of the channel (**Supp. Figure 3F**).

We next used this system to mechanically assess nuclei with perturbed Lamin A/C proteins. Control, HGPS, and *LMNA* KD fibroblasts were captured in the micro-mechanical device and compressed at four different strains until the channel was completely collapsed (**Figure 3C**). Confocal microscopy at each strain resolved fine differences in nuclear morphology during compression such that folds and wrinkles in the nuclear surface were observed to flatten as they contacted the channel walls (**Supp. Figure 4A**). HGPS and *LMNA* KD fibroblasts showed decreased resistance to nuclear lysis upon mechanical compression compared to control fibroblasts (**Figure 3D**). *LMNA* KD fibroblasts were particularly susceptible to lysis, which is in line with our observation that loss of Lamin A/C promotes nuclear rupture even in the absence of mechanical stress. In contrast, HGPS fibroblasts, which are less able to reorganize their nucleoscaffold in response to mechanical stress^60^, were able to resist lysis until the channel was completely closed.

To quantify nuclear deformation, ellipsoids were fitted to nuclei at each stage of compression and the elongation and flatness factors of each ellipsoid were calculated. Comparing these two factors during compression revealed a trend across all cell types in which nuclei transition from elongated morphologies towards flat, round morphologies (**Supp. Figure 4B,C**). To determine whether the device imposes mechanical stress on nuclei before the cells are captured in the channel, we assessed the morphologies of cells sitting in the inlet outside of the channel and those of cells sitting on glass coverslips (**Supp. Figure 4C**). We observed minimal differences between these populations. We further assessed each cell type’s ability to resist mechanical deformation by visualizing ellipsoid flatness and elongation as a function of channel width distal to the cell. Changes in fitted ellipsoid radii during compression showed that nuclear strain occurs both along and orthogonal to the channel, in contrast to the hydrogel beads (**Supp. Figure 4D**). HGPS and *LMNA* KD nuclei showed steeper increases in ellipsoid flatness with decreasing channel width compared to control nuclei, indicating reduced resistance to mechanical deformation (**Figure 3E**). These results were supported by atomic force microscopy (AFM) indentation experiments, which revealed significantly reduced Young’s moduli in HGPS and *LMNA* KD nuclei compared to control fibroblasts (**Figure 3F**). The maximum indentation depth was also greatest in HGPS nuclei, highlighting reduced mechanical stiffness (**Figure 3F**). These results are consistent with previous studies showing alterations in lamins confers changes in mechanical properties of nuclei^66^.

### Transient Nuclear Scaffold Alterations Accelerate Cell Fate Transitions

To assess whether the induced changes in nuclear morphology and chromatin organization with Lamin A/C modification impacted ability to adopt a new cellular fate, control, *LMNA* KD and HGPS fibroblasts were virally transduced to express reprogramming factors (OKSM)^67^ (**Figure 4A**). The number and size of reprogrammed stem cell colonies was assessed by TRA-1-60 live cell staining at 7, 15, 22 days after transduction and immunostaining for Oct3/4, Sox-2, and Nanog at 15 and 22 days (**Figure 4B,C**). HGPS cells failed to produce any sizable colonies over 22 days, but strongly expressed TRA-1-60, indicating robust initiation of the reprogramming process^68^. *LMNA* KD fibroblasts produced more TRA-1-60-positive colonies at days 15 and 22 when compared to controls, indicating accelerated reprogramming kinetics. To further assess the extent of reprogramming between conditions, we performed immunostaining for Oct3/4 and Nanog followed by flow cytometry (**Figure 4D-F**). We found comparable levels of Oct3/4 and Nanog at day 15 for *LMNA* KD and control fibroblasts, respectively, demonstrating a strong induction of the reprogramming program for both conditions (**Figure 4F**). We also observed a significant increase in Nanog expression for *LMNA* KD cells at day 22 compared to control fibroblasts, indicating an accelerated rate of genuine reprogramming. Immunofluorescence staining for Sox-2 and Nanog supported these findings whereby Sox-2 and Nanog expression were strongly activated in colonies that had formed by day 22 (**Supp. Figure 5**). HGPS fibroblasts were excluded from this analysis as the samples were too sparse to acquire reliable results. Overall, these results suggest that transient loss of Lamin A/C promotes accelerated colony formation with elevated *NANOG* expression, while permanently dysfunctional *LMNA* in HGPS cells promotes strong initiation of reprogramming that is hampered by complications from reduced proliferation and disorganized chromatin^69^.

**Figure 4.**
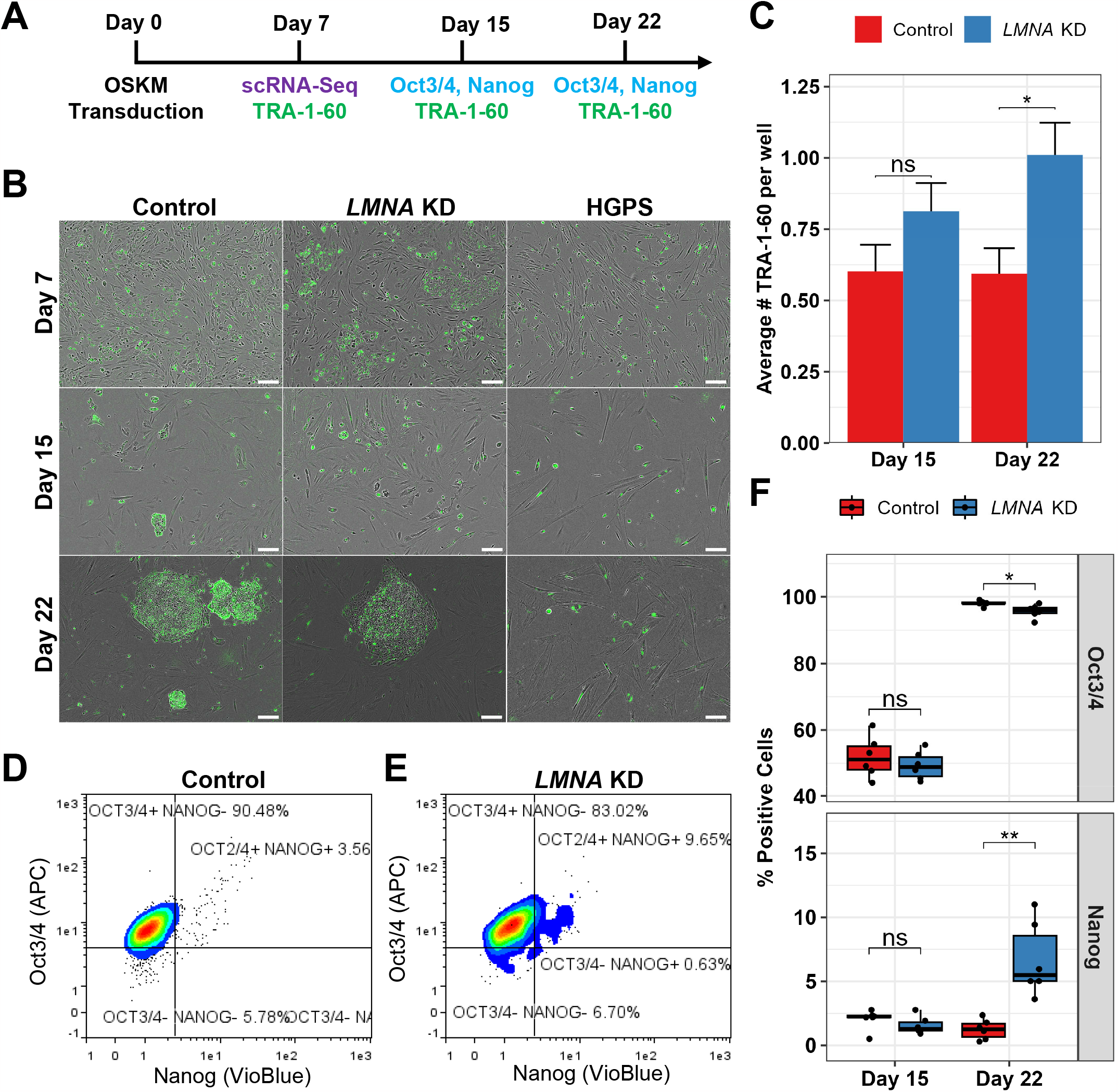
Altered Nuclear Morphology and Stiffness Converge to Modulate Reprogramming Kinetics. **A)** Overview of experimental design. **B)** Representative images of TRA-1-60 staining (green) overlaid with phase contrast images. Scale bars are 100 μm. **C)** Average number of TRA-1-60-positive colonies per well (Control, n=4; *LMNA* KD, n=6 wells). Representative flow cytometry plots for Oct3/4 and Nanog in **D)** control and **E)** *LMNA* knockdown fibroblasts 22 days after reprogramming. **F)** Quantitation of fibroblasts co-expressing Oct3/4 and Nanog at days 15 and 22 after reprogramming (n=6 wells each). Statistical comparisons are unpaired Mann-Whitney U-tests. *p<0.05, **p<0.01.

### Early Reprogramming Kinetics Are Altered By Loss of Functional Lamins

To further understand changes in the kinetics of early reprogramming from modifications to Lamin A/C, we generated and analyzed single-cell RNA sequencing datasets of control, HGPS, and *LMNA* knockdown fibroblasts at day 7 after reprogramming (**Figure 5, Supp. Figure 6A,B**). We identified a total of 7,893 cells (1,695 HGPS; 2,735 *LMNA* KD; 3,463 control), each expressing a mean of 27,815 unique molecular identifiers (UMIs) and 4,844 genes after basic quality filtering. Visualizing cell clustering in uniform manifold approximation and projection (UMAP) embeddings^70^ revealed transcriptionally distinct subpopulations that were common to all cell types (Cluster 2), enriched with HGPS fibroblasts (Cluster 6), or shared between control and *LMNA* KD fibroblasts (**Figure 5A**). Superimposing expression of pluripotency (*NANOG, GDF3*) and fibroblast lineage (*COL1A1, FN1*) genes onto UMAPs showed that non-reprogrammed cells abundantly expressed somatic genes distinct from cells undergoing reprogramming (**Figure 5B**). To characterize the progress of each cell type through reprogramming, we embedded the cells in a partition-based graph abstraction (PAGA) map^71^ and estimated diffusion pseudotime^72^ (**Figure 5C**). We identified two paths through the PAGA map that corresponded to two distinct reprogramming outcomes (**Figure 5D**). Path 1 (Cluster 3 → 1 → 4 → 0) captured cells that transitioned from expressing genes specific to the mesenchymal and fibroblast lineages towards expressing pluripotency genes. These same cells also expressed AP-1 factors at the beginning of the trajectory, which is consistent with our ATAC-Seq footprinting analysis (**Figure 2F**) and with these factors behaving as guardians of somatic cell fate^48,50,73^. Consistent with our *in vitro* characterizations, this path was also depleted for HGPS fibroblasts. Expression of the OSKM factors was enriched at the end of Path 1 and in Cluster 4, which was centrally located in the PAGA trajectories. Cells that followed Path 2 (Cluster 3 → 1 → 4 → 7 → 2) were predominantly HGPS cells that did not express pluripotency genes, but did express extracellular matrix remodeling genes characteristic of the fibroblast lineage and are known barriers to reprogramming^73^. The high expression of OSKM factors and the shared expression of fibroblast and pluripotent genes in Cluster 4 suggest that these cells represent a binary decision point between the different outcomes of Paths 1 and 2. We compared HGPS fibroblasts with control and *LMNA* KD fibroblasts in cluster 4 and found that while HGPS fibroblasts were expressing p53 programs related to senescence (*CDKN1A, CCNG1, DDB2*), the control and *LMNA* KD fibroblasts were enriched for genes related to cell proliferation, non-canonical Wnt signaling^74^ (*TCF7L1, WNT5A, FZD4*) and hedgehog signaling^75^ (*SEM1, SMOX, PSMA-4*, which are involved in embryonic development (**Supp. Figure 6C**). Moreover, these fibroblasts expressed genes related to multiple cell fates such as the neural (*ROBO3, SLIT3, SEMA5A*) and osteogenic (*COL1A1, SMAD1, TWIST2*) lineages, indicating the re-expression of previously silenced lineage specifiers. We further compared control and *LMNA* KD fibroblasts in Cluster 4 and found that *LMNA* KD fibroblasts upregulated anti-oxidant and Nodal signaling genes (*SQSTM1, DRAP1, PRDX6*), which are known to support pluripotency ^76,77^, while control fibroblasts expressed genes related to DNA methylation known to inhibit pluripotency induction (*GATAD2A*)^78^, peroxisome transport (*RAB8B*), and nuclear export of pluripotency genes (*SRSF5*)^79^ (**Figure 5E**). Summing these results shows that permanent modification of Lamin A/C, as observed in HGPS, reduces competency for changing cellular fate while transient loss of functional lamins modulates early reprogramming kinetics by upregulating unique pathways supportive of pluripotency.

**Figure 5.**
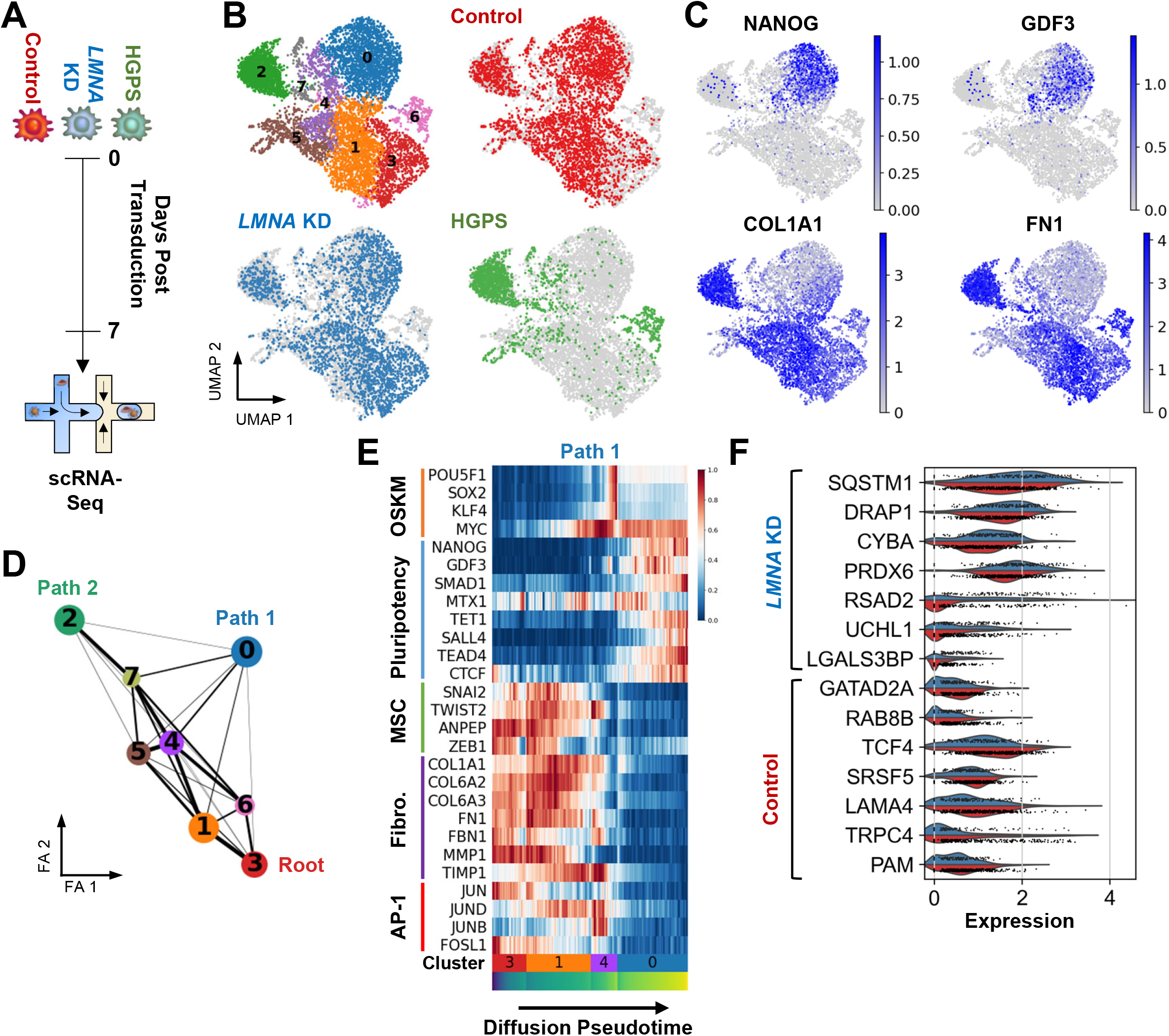
Early Reprogramming Cells with Varying Nucleo-Scaffold Composition Transition Along Distinct Reprogramming Trajectories. **A)** Schematic of transduction of fibroblasts from three conditions with reprogramming vector and isolation 7 days followed by profiling with single-cell RNA sequencing (scRNA-Seq). **B)** UMAP embeddings colored by Leiden clustering and each sample. **C)** UMAP embeddings showing the expression of pluripotency (top row) and fibroblast (bottom row) genes. **D)** Partition-based graph abstraction (PAGA) analysis of inferred trajectories in early reprogramming cells. Distinct trajectories emerging from the root node (Cluster 3) are shown. **E)** Reconstructing gene expression changes of mesenchymal (MSC) and fibrogenic (Fibro.) fates transitioning towards distinct reprogramming outcomes along PAGA paths as a function of diffusion pseudotime. Gene expression is normalized and scaled to unit values across each row within each path. **F)** Split violin plot of differentially expressed genes between control and *LMNA* knockdown fibroblasts in Cluster 4. Genes upregulated in each condition are marked at the bottom.

## DISCUSSION

Cellular identity is regulated by the coupled actions of lineage-specific TF binding, and a chromatin landscape that represses genes associated with alternate fates^2^. Here we show that modifying Lamin A/C changes nuclear morphology and elasticity^80,16^, and opens condensed chromatin to facilitate alterations in the ability to change cell fate^12,18^. Recently, mechano-transductive strategies^81^ have been shown to enhance the efficiency of the reprogramming process through reduction of heterochromatin^82,83^, which is a well-known barrier to reprogramming^84^ and contributes to proper expression of lineage-specific programs^85,86^. Our results show that temporary perturbation of the nuclear scaffold from *LMNA* KD can transiently increase constitutive heterochromatin markers such as H3K9me3,^85,86^ as well as increase chromatin accessibility in constitutive LADs. These chromatin changes occurred around genes related to PIWI RNA metabolism, which are known to repress transposable elements that are typically silenced within LADs. Thus, while H3K9me3 may be temporarily increased in *LMNA* KD cells, lineage-specific genes that are typically repressed within LADs may also be opening. This result indicates that the kinetics of reprogramming somatic cells back to pluripotency are mediated by heterochromatin detachment within LADs, which is in line with previous improvements in reprogramming efficiency by treatment with vitamin C^87^, which specifically targets heterochromatin at the nuclear periphery^88^.

Hutchinson-Gilford Progeria Syndrome is a devastating rare condition in which functional Lamin A/C is replaced by mutant progerin. Replacement of Lamin A/C with progerin contributes to aberrant nuclear morphology, loss of H3K9me3, and opening of genes normally repressed in LADs^34,89,90^ suggestive of an increased propensity to change cellular fate. Our results show reductions in the ability of HGPS cells to reprogram back to pluripotency and attribute this to reductions in proliferative capacity from accumulation of progerin^90^. Cellular reprogramming to pluripotency requires concerted proliferation and multiple passages of HGPS cells^91^ have been shown to contribute to senescence^92^. Single-cell RNA sequencing of fibroblasts early in the reprogramming process further substantiate this effect and showed that while control and *LMNA* KD fibroblasts traveled along similar trajectories towards pluripotency, HGPS fibroblasts did not appreciably express pluripotency genes and varied in their trajectory. These results suggest that temporary changes in lamina-associated heterochromatin facilitate formation of new chromatin accessibility patterns that are permissive to cell fate changes, but permanent LAD detachment and progerin accumulation contributes to dysfunctional cell cycle regulation and senescence programs^93^.

Mammalian cells are sensitive to tissue stiffness^13^ and adjust the mechanical properties of their nuclei by modifying expression of lamin isoforms and re-distributing heterochromatin. Techniques to connect biophysical properties of nuclei with morphological changes have been lacking^94^. Our microfluidic squeezing approach begins to address this challenge by facile cell trapping in optically transparent microfluidic channels and controllably deforming nuclei. Monitoring the squeezing process with confocal microscopy facilitated observation of increases in nuclear deformation for HGPS and *LMNA* KD fibroblasts when compared to controls, which mirrored results obtained with AFM indentation. Given this system is adaptable to integrate with light-sheet microscopy^95^ for tracking of faster cellular processes, is scalable to probe deformation of individual nuclei in multinucleated cells such as muscle, and displayed sensitivity to follow the unfolding of nuclear wrinkles, which can be used to predict mechano-transduction ability^96^ and resistance to nuclear lysis; we envision additional aspects of nuclear mechanobiology can be accessed by this novel approach. Our observations revealed HGPS and *LMNA* KD fibroblasts displayed increased propensity to lyse under smaller nuclear strains, and we speculate that cells in mechanically active tissues such as the myocardium, skeletal muscle and vasculature may attempt to remodel their matrix as a protective mechanism. In line with this, recent evidence suggests that cell types in mechanically active tissues with laminopathies exhibit pathological symptoms at faster rates than tissues with less deformation^97^.

The biophysical changes in nuclear structure that are imparted from modifications to lamins present an exciting opportunity to further understand links between chromatin organization and epigenetic factors that enforce lineage-specific gene expression^69,98,99^. Future work in this area will focus on how changes to the nuclear scaffold impact DNA interactions with other nuclear bodies and self-interacting topological associated domains.

## Materials and Methods

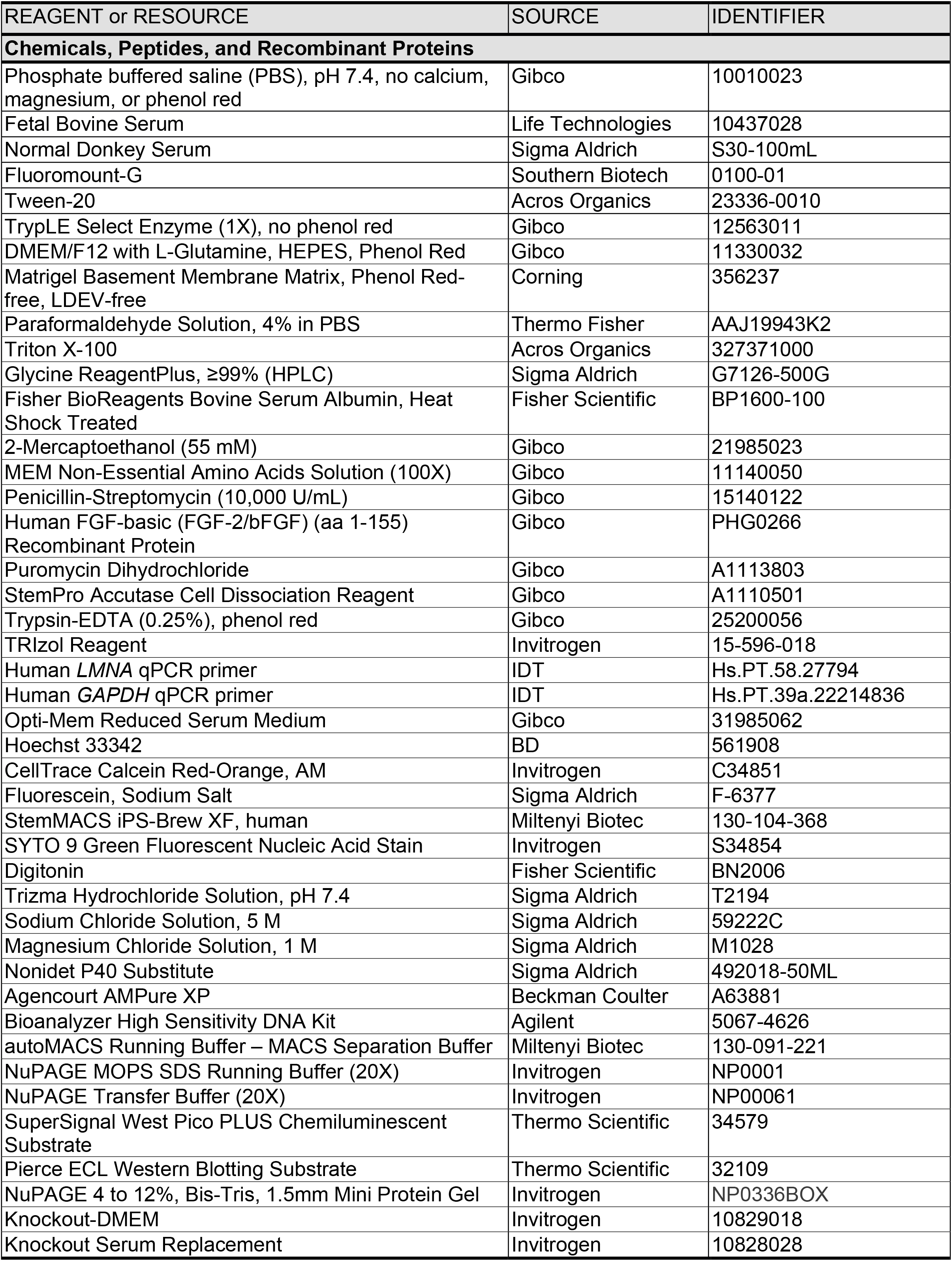
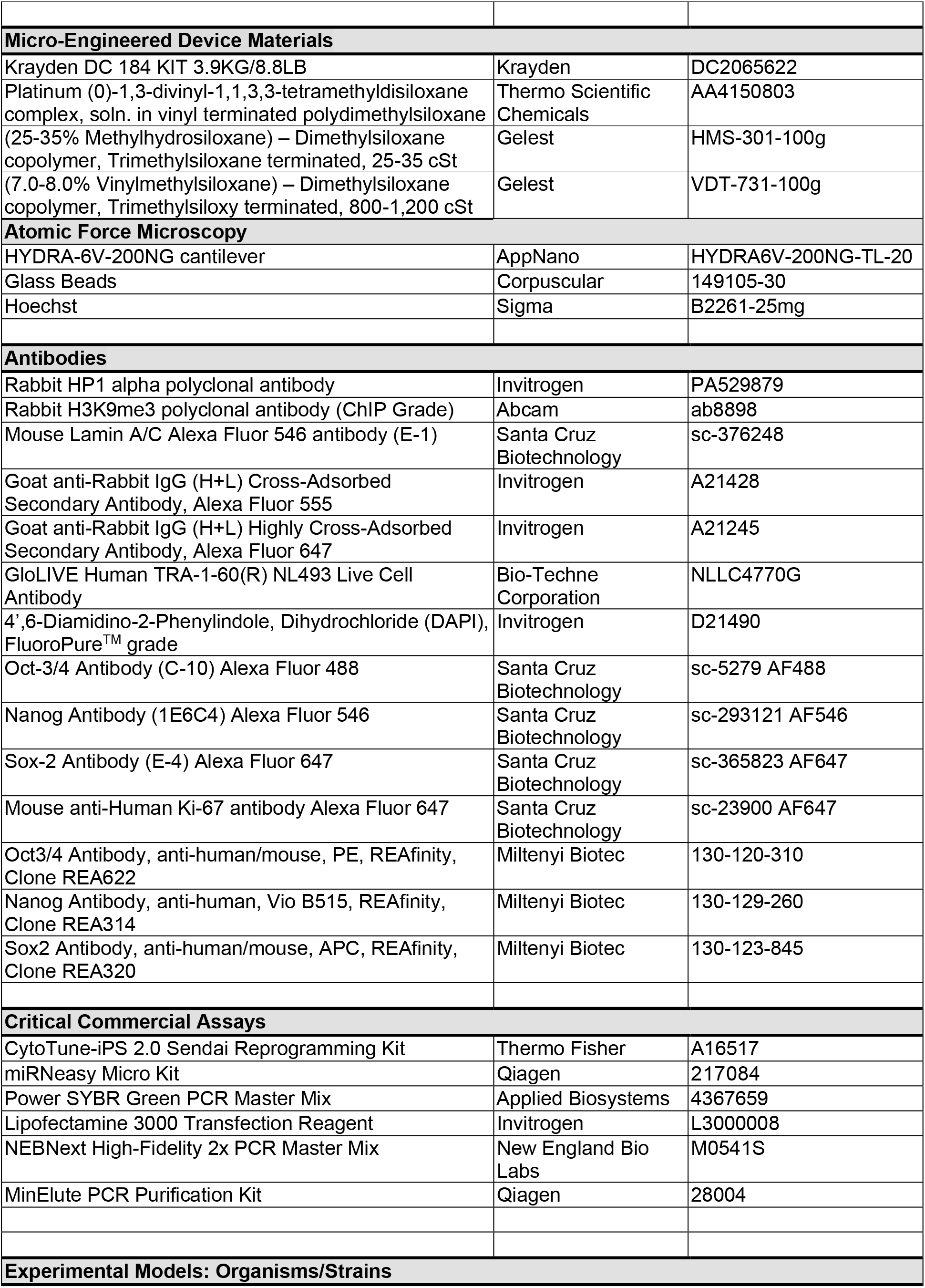
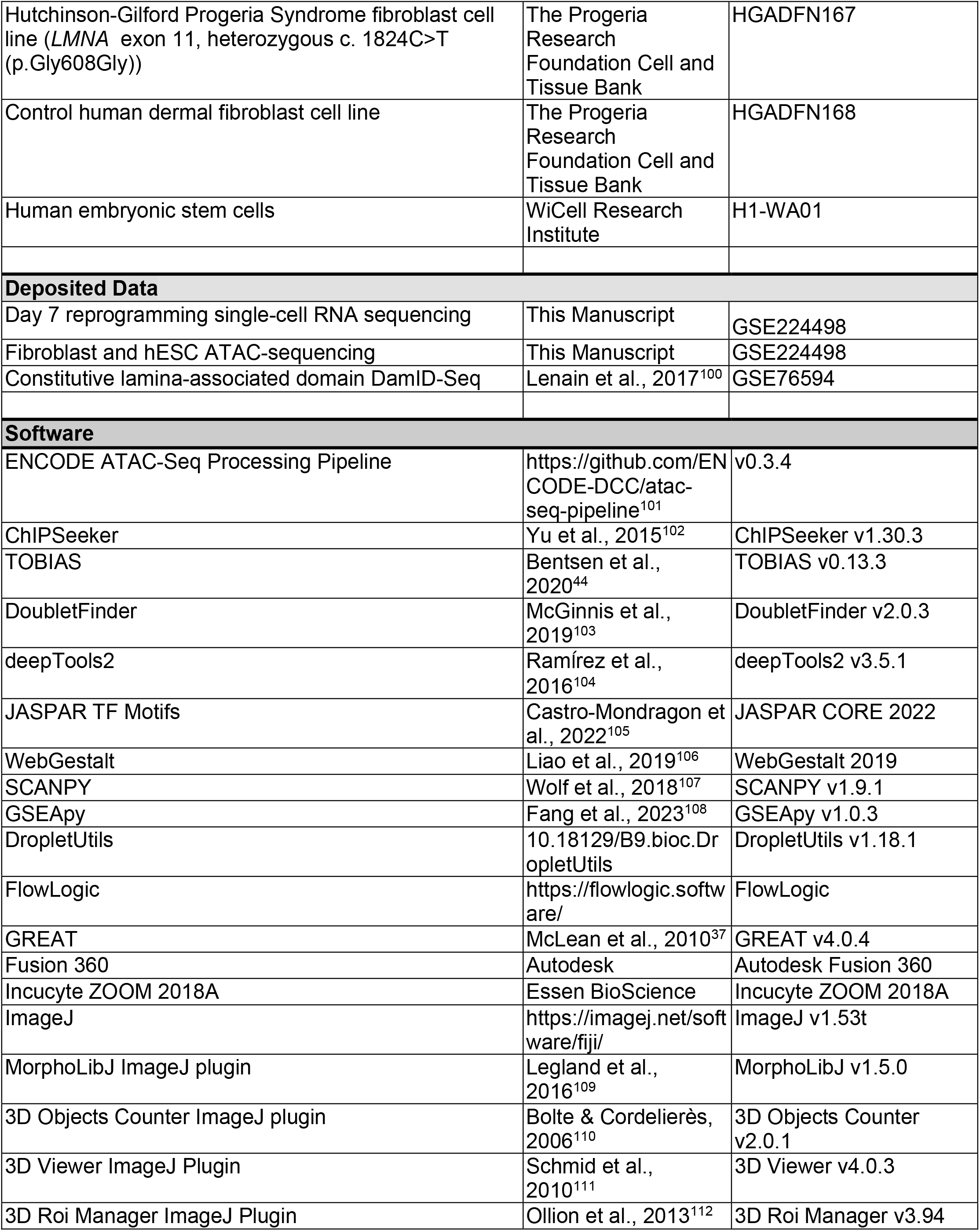

### Cell culture

Dermal fibroblasts from a normal adult and a patient with Hutchinson-Gilford Progeria Syndrome (HGPS) were obtained from The Progeria Research Foundation Cell and Tissue Bank. Prior to seeding cells, culture plates were coated in a 1:30 dilution of Matrigel (Corning) in DMEM/F12 for 1 hour at room temperature.Extra plates were kept at 4C for at most 2 weeks and allowed to equilibrate for 1 hour at room temperature before use. The growth media consisted of DMEM/F12 (Gibco), 10% FBS, 0.1 mmol/L nonessential amino acids (Gibco), 1 mmol/L L-glutamine (Gibco), 0.1 mmol/L beta-mercaptoethanol, and 1X penicillin-streptomycin. Fibroblasts were passed at ∼80% confluency using 0.25% Trypsin-EDTA (Gibco).

Human embryonic stem cells were grown on ESC-qualified Matrigel (BD) in human ES media consisting of KO-DMEM (Invitrogen), 10% KOSR (Invitrogen), 1% glutamax or L-glutamin, NEAA (Thermo), penicillin/streptomycin, 0.1% 2-mercaptoethanol and 10 ng/mL human bFGF (Gibco). All cells were passaged using Accutase (Invitrogen).

### siRNA transfection and real-time PCR analysis

#### LMNA knockdown in human dermal fibroblasts

One day prior to transfection, human dermal fibroblasts were seeded in 12-well or 24-well Matrigel-coated plates at 15,000 or 10,000 cells per well, respectively. Cells were seeded on glass coverslips for immunostaining. On the day of transfection, cells were treated with 50 nM of DsiRNA against Lamin A/C (IDT; hs.Ri.Lmna.13.3) using Lipofectamine 3000 (Thermo Fisher) and allowed to recover for 5 days. The medium was replaced three days after transfection.

#### Quantitative PCR

Titration of *LMNA* DsiRNAs and knockdown efficiency were confirmed in part using quantitative PCR. Cells were harvested using Trizol, and a phenol-chloroform RNA extraction was performed using the Qiagen miRNeasy Micro Kit according to the manufacturer’s protocol. cDNA was then synthesized using the Superscript III First-Strand Synthesis kit and quantitative PCR (qPCR) was performed using Power SYBR Green PCR mix (Thermo Fisher) on a QuantStudio 3 thermocycler (Applied BioSystems). The *LMNA* primer was purchase from IDT (PrimeTime qPCR assay, Hs.PT.58.27794).

#### Proliferation assessment

To confirm that lipid nanoparticle transfection has minimal effect on cell proliferation, we compared the proliferation of control and *LMNA* knockdown cells using the BioSpa Live Cell Analysis system (BioTek) and performed Ki67 immunofluorescence staining (Santa Cruz Biotechnology).

### Cell immunostaining and analysis

#### Immunostaining

Cells were fixed using 3% paraformaldehyde in PBS for 10 minutes and permeabilized with 0.1% Triton X-100 for 10 minutes at room temperature. The cells were then washed and blocked using a solution of 0.005% Triton X-100 and 0.1% normal donkey serum in PBS for 30 minutes at room temperature. Primary antibodies were added in the same buffer for 1 hour at room temperature in the dark followed by secondary antibodies using the same procedure. DAPI stain was diluted to 1.5 ug/mL in PBS, and the cells were incubated with diluted DAPI for 1 minute at room temperature, in the dark. Cells cultured on glass coverslips were then lifted and mounted on microscope slides using Fluoromount-G Mounting Medium (Southern Biotech) and stored at 4C overnight. The primary antibodies used in this study were: Rabbit anti-HP1a (Invitrogen, PA529879), Rabbit anti-H3K9me3 (Abcam, ab8898), and Mouse anti-Lamin A/C-Alexa Fluor 546 (Santa Cruz Biotechnology, sc-376248). Secondary antibodies were Goat anti-Rabbit Alexa Fluor 555 or 647 (Fisher Scientific).

#### Fluorescence imaging

Fluorescence images were acquired using a Zeiss AxioVert epifluorescent microscope at 40X magnification or a Nikon A1R inverted confocal microscope at 60X. The same acquisition parameters were used across samples for quantitative microscopy. All image processing was done in ImageJ. Nuclei were segmented using the DAPI stain and used to quantify the integrated fluorescence intensity of the other stains^113^. Varying segmentation and image processing protocols were implemented depending on the objective magnification and sample type. Background regions of interest (ROI) were drawn by hand in each image and used to calculate corrected total cell fluorescence (CTCF):

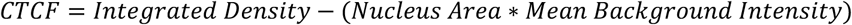

### Nuclei morphology analysis

Image analyses were performed in ImageJ and processed using R (v4.1.3). Cells for each condition were fixed and immunostained for Lamin A/C. The Huang threshold algorithm was used to segment and threshold nuclei in the corresponding Lamin A/C channel to produce a binary image. A pixelated ROI boundary was acquired for each nucleus using the wand tool and smoothed by sub-pixel interpolation with 1 pixel resolution. The resulting cartesian coordinates were saved for further analysis. For each nucleus, a circle with the same centroid and area was constructed, and the ratio between the centroid-to-perimeter distance and the radius of the equivalent circle was calculated as its relative radius. To reduce noisiness in the data, a cubic smoothing spline was applied using the R stats package with a smoothing parameter of 0.45. Smoothed relative radii were graphed as a function of degrees in a clockwise direction. Invaginations and protrusions in the nuclear envelope were quantified by identifying local minima and maxima of the smoothed profiles, respectively, with a sliding window of 10 degrees. The average number of invaginations and protrusions for each LINC knockdown were compared against that of wild-type nuclei using Mann-Whitney U-tests.

### Western blots

Cell pellets from control and HGPS fibroblasts were resuspended in 2x Laemmli buffer and vortexed. Equal volumes of lysates were run on NuPAGE 4-12% Bis-Tris 1.5 mm protein gels (Invitrogen) in MOPS SDS buffer (Invitrogen) for 2 hours at 80 V then transferred to nitrocellulose paper. The resulting membranes were cut to isolate the proteins of interest. Lamin A/C and progerin were located around 70 kDa while GAPDH was located around 40 kDa. Membranes were then washed three times in PBS-T (1mL Tween 20 in 1L 1X PBS) for five minutes each and blocked with 5% milk in PBS-T at room temperature for 30 minutes. Primary antibodies were then incubated on each membrane in 5% milk in PBS-T overnight at 4C (1:1000 GAPDH, 1:200 Lamin A/C). The following day, membranes were washed five times in PBS-T and incubated with HRP secondary antibody (1:500 Lamin A/C, 1:200 GAPDH). After washing, the membranes were imaged on a Chemi-Doc using the Pierce ECL kit (Thermo Fisher) for GAPDH and the SuperSignal Pico PLUS kit (Thermo Fisher) for Lamin A/C. Blot intensities for GAPDH were measured using the ImageLab software (Bio-Rad) and used to normalize sample volumes. The protocol was repeated with these normalized sample volumes for quantification.

### Microfluidic assessment of nuclear mechanical properties

#### Fabrication of micro-engineered devices

We utilized deterministic cracking^114^ of thin polymeric films at patterned sites to generate fluidic channels that are deformable under uniaxial tension and to trap single cells^115^. First, we used photolithography to create a mold with two horizontal arrays of v-shaped grooves that were positioned across from each other with the tips of each “v” facing inwards. Each pair of grooves defined the positions of a single crack and served as inlets and outlets into the cracked channel. Hard polydimethylsiloxane (h-PDMS), which is composed of vinyl and hydrosilane end-linked polymers^116^ that increase its Young’s modulus and render the material more brittle, was poured over the mold and spun on a Brewer Science Spin Coater to create a thin film (Step 1 – 500 RPM, 5 seconds, 250 RPM/sec ramp; Step 2 – 2000 RPM, 40 seconds, 250 RPM/sec ramp). The mold was then incubated in a 120 C oven for 2 minutes and allowed to cool for 30 seconds at room temperature. 20 g of 1:10 PDMS was then poured over the h-PDMS film and incubated in a 60 C oven for at least 4 hours. The mismatch in moduli between the h-PDMS film and PDMS substrate promoted crack formation between each pair of grooves^64,117^. Channels were created by applying uniaxial tension across the resulting device using a custom fabricated clamp integrated to a micrometer displacement stage. Each device was observed under a dissection microscope during tension to monitor crack formation. Inlets and outlets were then created with a 0.75mm punch. To form the bottom wall of the channels, a thin film of 1:10 PDMS was spun onto glass slides (Step 1 – 500 RPM, 5 seconds, 250 RPM/sec ramp; Step 2 – 1000 RPM, 60 seconds, 250 RPM/sec ramp) and bonded to the h-PDMS/PDMS device using a plasma asher (Femto Science).

#### Measurement of bulk PDMS and h-PDMS stiffness

The Young’s moduli of the bulk PDMS and h-PDMS layers in the device were measured under strain-controlled uniaxial tension using a Dynamic Mechanical Analyzer (TA Instruments, RSA III). Samples were cast into a 3-D printed mold following the ASTM D638 Type 1 standard. Samples were stretched at 0.01 mm/s for 10 seconds and normal stress was measured at 300 time points assuming a temperature of 21°C.

#### Production of alginate hydrogel beads

Alginate hydrogels with a 2.4 kPa Young’s modulus were generated as previously described^65^. Briefly, a gel solution of DMEM and 2% w/v lyophilized alginate conjugated with Rhodamine B was encapsulated in microfluidic water-in-oil droplets with an outer oil phase comprising 1% acetic acid in PFPE surfactant. A secondary flow of CaCO_3_ solution in DMEM was added to form alginate hydrogel emulsions that were filtered through a 40 μm cell strainer.

#### Loading and squeezing samples in the device

Each device was washed with 70% ethanol and 1X PBS prior to sample loading. Devices were stretched in the custom clamp, which opened the channel, and a pair of pipettes generated a negative pressure gradient across the length of the channel by dispensing cleaning solution in the inlet and aspirating the solution through the outlet. Cells and hydrogel bead samples were loaded in the same manner. To trap samples, the strain on the device was relaxed to constrict the channel. The applied strain was recorded at rest before the channel was opened and at the moment of sample capture to monitor channel dimensions. To compress samples, uniaxial strain on the device in the clamp was incrementally removed in 0.5mm steps until the channel was completely collapsed.

#### Image acquisition

All confocal images were acquired on a Nikon A1R confocal microscope with a 20X air objective. After the channel cleaning and prior to sample loading, the channel was filled with an aqueous solution of fluorescein to visualize channel dimensions. To image cells, the channel was filled with a solution of fluorescein in PBS without calcium or magnesium. To image alginate hydrogel beads, the channel was filled with a solution of fluorescein in DMEM with ∼1.8mM calcium chloride to minimize swelling. Prior to loading, cells were stained with Calcein AM Red-Orange and Hoechst 33342, and resuspended at 0.5×10^6 cells/mL in complete growth media. Cells and hydrogel bead suspensions were kept on ice during image acquisition. Confocal z-stacks (1 μm z-steps) were acquired after cell trapping and at each strain during compression.

#### Image analysis

All image analysis done in ImageJ. Briefly, samples were segmented from z-stack images using the MorphoLibJ^109^ (https://imagej.net/plugins/morpholibj) and 3D Objects Counter^110^ (https://imagej.net/plugins/3d-objects-counter) plugins. This allows surface area and volume measurements to be recorded. Next, the thickness of each sample in orthogonal directions is recorded in addition to measurements of the orthogonal cross-sections (e.g. aspect ratio, area, perimeter). Finally, an ellipsoid is fitted to each sample using the 3D ROI Manager^112^ (https://imagejdocu.list.lu/plugin/stacks/3d_ij_suite/start). Three-dimensional visualizations of the channel and samples during compression were generated using the 3D Viewer plugin^111^ in ImageJ without resampling. STL files of the bead and channel surfaces were exported from 3D Viewer and opened in Fusion 360 for 3D renderings. The imported meshes were adaptively re-meshed using density and shape preservation settings of 3 and 1, respectively, and rendered images were captured using the orthographic camera setting.

### Atomic Force Microscopy

To measure nuclei stiffness, atomic force microscopy (AFM) nanoindentation in contact mode was performed using a Nanosurf FlexBio atomic force microscope with a HYDRA6V-200NG (AppNano) probe tip (k=0.0348 N/m) affixed with an 11.39 μm diameter glass microsphere (Fisher). Cells were cultured on 18 mm glass coverslips and stained with Hoechst dye (1:1000) to visualize nuclei location. The glass coverslips were then adhered to a 50 × 55 mm glass coverslip that was then placed onto the AFM stage. The probe tip was equilibrated in the cell media above the culture monolayer for 20 min prior to indentation measurements. Indentations were performed on 15 nuclei per condition. Force-displacement curves were fit to the Hertz model assuming a Poisson’s ratio of 0.5 to determine Young’s Modulus values.

### Omni-ATAC sequencing and analysis

#### Library generation

10-20,000 human dermal fibroblasts before and after *LMNA* knockdown and human embryonic stem cells were centrifuged at 500xg for 5 min at 4°C in a fixed angle centrifuge. After removing the supernatant, the cell pellets were re-suspended in 50uL ATAC-Resuspension Buffer (RSB) (500uL 1M Tris-HCl, 100uL 5M NaCl, 150uL 1M MgCl_2_ in 49.25mL sterile water) with 0.1% NP40, 0.1% Tween-20, and 0.01% Digitonin. Samples were pipetted up and down three times to mix, incubated 3 minutes on ice, then diluted in 1mL of ice-cold ATAC-resuspension buffer (RSB) containing 0.1% Tween-20 and centrifuged at 500xg for 10 min at 4°C to pellet the nuclei. The supernatants were carefully removed and the pellets were re-suspended in 25 μL of transposase reaction mix (12.5 μL 2×TD buffer, 1.25 μL TN5 transposase (Illumina) and 11.25 μL nuclease-free water). Transposase reactions were carried out by incubating samples at 37°C for 30 min under mild agitation (300 rpm on a Thermo-mixer C, Eppendorf). Once the incubation was completed,sample tubes were placed on ice and the transposed DNA fragments from each sample were purified using a Qiagen MinElute PCR Purification Kit following manufacture’s protocol. Purified DNA fragments were then amplified for 13 cycles using barcoded PCR primers and NEB Next High Fidelity 2× PCR Master Mix (New England Bio Labs) on a thermal cycler. Double concentrated Ampure beads were used to purify transposed DNA amplicons. The molarity of each DNA library was determined (Agilent 2100 Bioanalyzer), pooled into a single tube and sequenced on a NextSeq (Illumina) using 76-bp paired-end reads.

#### Library pre-processing

The ATAC-Seq samples were analyzed with the ENCODE ATAC-seq processing pipeline (https://github.com/ENCODE-DCC/atac-seq-pipeline, version 0.3.4)^101^. The naive overlap peak set from all replicates for a given sample was used. Such peak sets from all samples were concatenated and merged using the bedtools merge command to produce a master peak set across all samples.

#### Peak annotation, motif enrichment, and gene set enrichment

For principal component analysis (PCA), the merged peak set was truncated to 200 bp regions surrounding peak summits and fragments counted using bedtools coverage. The resulting matrix was normalized using the limma package^118^ before PCA. Heatmaps of shared and unique sites were generated by assigning the nearest TSS to each site and calculating chromatin accessibility in a 2 kb region surrounding each TSS using the computeMatrix function in the deepTools package^104^. Accessible sites were annotated for genomic regions using the ChIPseeker^102^ package in R with a TSS region of +/- 1 kb. The resulting annotations were collapsed into promoter, genic, and distal regions. Gene ontology (GO) enrichment of accessible sites was performed using the GREAT algorithm^37^ with the hg38 genome and the basal plus extension method for assigning genomic regions to genes (5 kb proximal; 1 kb downstream; distal up to 10 kb). Enriched GO terms were identified with hypergeometric fold enrichments greater than 2 and hypergeometric FDR values less than 0.05. The foreground for this analysis was unique sites for each condition and the background set was the merged peak set. This identified terms that were uniquely enriched in each condition^119^.

#### Transcription factor footprint analysis

TF motifs were downloaded from the JASPAR CORE 2022 database^105^ and used for footprint analyses with the TOBIAS toolkit (v0.13.3)^44^. The bam mm10 alignments from the ENCODE ATAC-seq pipeline were corrected for Tn5 enzymatic bias using the ‘ATACorrect’ function. Blacklisted regions^120^ were excluded from the bias estimations. To perform differential TF binding analysis, a union set of ATAC peaks was generated across control and *LMNA* knockdown fibroblasts using the bedtools merge command. Continuous footprinting scores for the bias-corrected ATAC signals were then calculated across the union peak set using the ‘ScoreBigwig’ function. Differential testing of TF binding was calculated using the ‘BINDetect’ function and significant TF binding events were identified as those above the 95^th^ percentile or below the 5^th^ percentile of differential binding scores or those above the 95^th^ percentile of -log10(p-values).

#### Chromatin accessibility in constitutive lamina-associated domains

DamID maps and coordinates of lamina-associated domains (LADs) from cycling embryonic human lung fibroblasts (Tig3) were downloaded from the GEO Accession Database (GSE76594)^100^. ATAC signals across samples were summed in and plotted across constitutive LAD domains using the computeMatrix and plotProfile functions in the deepTools package^104^.

### Assessments of somatic reprogramming kinetics

#### Somatic cell reprogramming

Reprogramming efficiency with OSKM factors^67^ was compared between human dermal fibroblasts from normal adults and HGPS patients, and human dermal fibroblasts after *LMNA* knockdown. Control and HGPS fibroblasts were cultured in Matrigel-coated 6-well plates and passaged twice before reprogramming. *LMNA* knockdown was performed in control fibroblasts as previously described and all cells were re-seeded at 45,000 cells/well in separate Matrigel-coated 12-well plates two days before reprogramming. On the day of reprogramming, one well of each plate was lifted for multiplicity of infection (MOI) calculations. 4 wells of each plate were transduced with the CytoTune-iPS 2.0 Sendai Reprogramming Kit (Thermo Fisher) using the recommended MOIs for each factor (KOS=5, hc-Myc=5, hKlf4=3) in fibroblast growth media. Media was exchanged every other day for seven days. On the seventh day, 10,000 cells from each well was seeded into three or four Matrigel-coated 12-well culture plates and fibroblast growth media was replaced with StemMACS iPS-Brew XF stem cell culture media (Miltenyi Biotec) and changed every day. A portion of each sample was seeded on Matrigel-coated glass coverslips for immunofluorescence imaging. All cells were cultured at 37C with 5% CO2.

#### Live TRA-1-60 staining and analysis

All samples were stained with 1X GloLIVE Human TRA-1-60 NL493 Live Cell antibody (R&D Systems) in stem cell media for 30 minutes at 37C on days 7, 15, and 22 after reprogramming. Media was then replaced and the cells were imaged at 37C using the Incucyte ZOOM (Essen Bioscience) with 16 images per well at 4X magnification. Phase contrast and GFP filter images were acquired. Image analysis was performed in ImageJ. Masks for the phase contrast and GFP images were generated and the average TRA-1-60 confluence was calculated as the ratio of total areas per image. The number of TRA-1-60-positive colonies was counted by hand.

#### Immunostaining

Immunofluorescence staining was performed at days 15 and 22 as previously described. Primary antibodies were all acquired from Santa Cruz Biotechnology and used at 1:100 dilution: Nanog (sc-293121 AF546), Oct3/4 (sc-5279 AF488), and Sox-2 (sc-365823 AF647). All samples were counterstained with DAPI and imaged on either the Nikon A1R inverted confocal microscope at 60X or the Zeiss AxioVert epifluorescent microscope at 10X.

#### Single-cell RNA sequencing and analysis

7 days after reprogramming, control, Lamin A/C knockdown, and HGPS fibroblasts were each labeled with 3’ CellPlex cell multiplexing oligos (CMO; 10X Genomics) and pooled together before loading onto the 10X Genomics chromium single cell controller for single-cell RNA sequencing (Single Cell 3’ v3 chemistry). The CMOs used for each sample were: Control-CMO309, HGPS-CMO310, and *LMNA* KD-CMO311. 30,000 total cells were targeted with 50,000 reads per cell. Libraries were sequenced on a NovaSeq 6000 (Illumina) and aligned to the refdata-gex-GRCh38-2020-A reference transcriptome using CellRanger multi with default parameters (v7.1.0; 10X Genomics). Samples were demultiplexed using the hashedDrops function in the DropletUtils package (v1.18.1). Demultiplexed cells were identified with an FDR threshold of 1e-5 and a minimum log fold change confidence level in CMO assignment of 2. Confidently assigned cells were processed using the SCANPY package (v1.9.1)^107^. Cells with more than 250 genes, 500 UMIs, and less than 8% mitochondrial reads were retained for downstream analyses. Genes expressed in fewer than 3 cells were also removed. Counts were normalized across cells and scaled to 1e4 per cell and highly variable genes were identified. Effects due to total UMIs and mitochondrial reads per cell were regressed out and the resulting counts were scaled to unit variance with maximum variance clipped at 10. Cell cycle scoring showed minimal relationship between cell cycle phase and count data. A neighborhood graph was constructed in PCA space followed by Leiden clustering with a resolution of 0.4. Next, partition-based graph abstraction (PAGA)^71^ analysis was performed in the SCANPY package and used to initialize all UMAP embeddings. Gene expression overlays on UMAP embeddings were clipped at the top 99^th^ percentile. Marker genes were identified between clusters using the Wilcoxon test (padj<0.01) and annotated with GO and Reactome terms (padj<0.01) using EnrichR^121^ through the GSEApy package (v1.0.3)^108^. Finally, diffusion pseudotime trajectory analysis was performed using the Force Atlas 2 graph layout algorithm^122^ with PAGA initialization. Genes that varied over diffusion pseudotime were visualized using the paga_path command in SCANPY with the expression values per gene normalized and scaled to 1.

#### Flow cytometry

Flow cytometry of Nanog, Oct3/4, and Sox-2 was performed using the MACSQuant VYB (Miltenyi Biotec) on days 15 and 22 after reprogramming. Due to limited cell availability, 1-2 wells of HGPS fibroblasts were assayed at each time point while 6 wells were assayed for control and *LMNA* knockdown fibroblasts. Cells were lifted in 1X TrypLE Select and pelleted in 1.5 mL Eppendorf tubes. The supernatant was aspirated and the pellet was resuspended in 200 μL cold 3% paraformaldehyde in PBS for 10 minutes on ice. 1 mL of cold 0.1% Tween-20 in PBS with 3 mg/mL bovine serum albumin (BSA) was then added and incubated on ice for 15 minutes. The samples were then spun down and the supernatant removed. The pellets were resuspended in 50 μL of pooled REAfinity antibodies (Miltenyi Biotec, 1:50 dilution) for 40 minutes on ice. The antibodies used were Anti-human/mouse *OCT3/4*-PE, Anti-human *NANOG*-Vio B515, and Anti-human/mouse *SOX2*-APC. Nuclei were counterstained with 5 μM of SYTO 9 Green (Invitrogen) for 10 minutes on ice. Due to cell availability, only one negative control was provided for all samples. Single-stained controls were also included for gating. Analysis was performed using the FlowLogic software (https://flowlogic.software/).

## Acknowledgments

The authors thank the University of Michigan DNA Sequencing Core for assistance with sequencing. The authors also thank members of the Takayama and Aguilar laboratories, especially Krithika Balakrishnan and Kanishka da Silva. Research reported in this publication was supported by a National Science Foundation CAREER award (2045977), the National Institute of Arthritis and Musculoskeletal and Skin Diseases of the National Institutes of Health under Award Number P30 AR069620 (C.A.A.), the National Institute of General Medical Sciences of the National Institutes of Health under Award Number R01 GM123517 (S.T., M.D.T.), the 3M Foundation (C.A.A.), American Federation for Aging Research Grant for Junior Faculty (C.A.A.), the University of Michigan Geriatrics Center and National Institute of Aging under award number P30 AG024824 (C.A.A.), the Department of Defense and Congressionally Directed Medical Research Program W81XWH2010336 and W81XWH2110491 (C.A.A.), Defense Advanced Research Projects Agency (DARPA) “BETR” award D20AC0002 (C.A.A.) awarded by the U.S. Department of the Interior (DOI), Interior Business Center, and the National Institute of Biomedical Imaging and Bioengineering Training Award T32 EB005582 (BAY). The content is solely the responsibility of the authors and does not necessarily represent the official views of the National Institutes of Health.

## Accession Code

GSE224498

## Author contributions

B.A.Y. performed the experiments and analyzed the data. I.N. and P.M.F. conducted the initial proof of concept experiments. H.H. performed the atomic force microscopy experiments. A.M.R. contributed to the design and execution of the reprogramming experiments. S.W. produced the alginate hydrogels. J.L. contributed to the ATAC-Seq library preparation and A.S. contributed to the ATAC-Seq data analysis. S.T., B.M.B., J.W.S., and M.D.T. contributed reagents and tools. C.A.A. directed the project, designed the experiments, and guided the analysis. M.D.T designed the mechanics experiments and guided the analysis. B.A.Y. and C.A.A. wrote the draft, and all authors contributed to the editing of the manuscript.

---

**Supplemental Figure 1.**
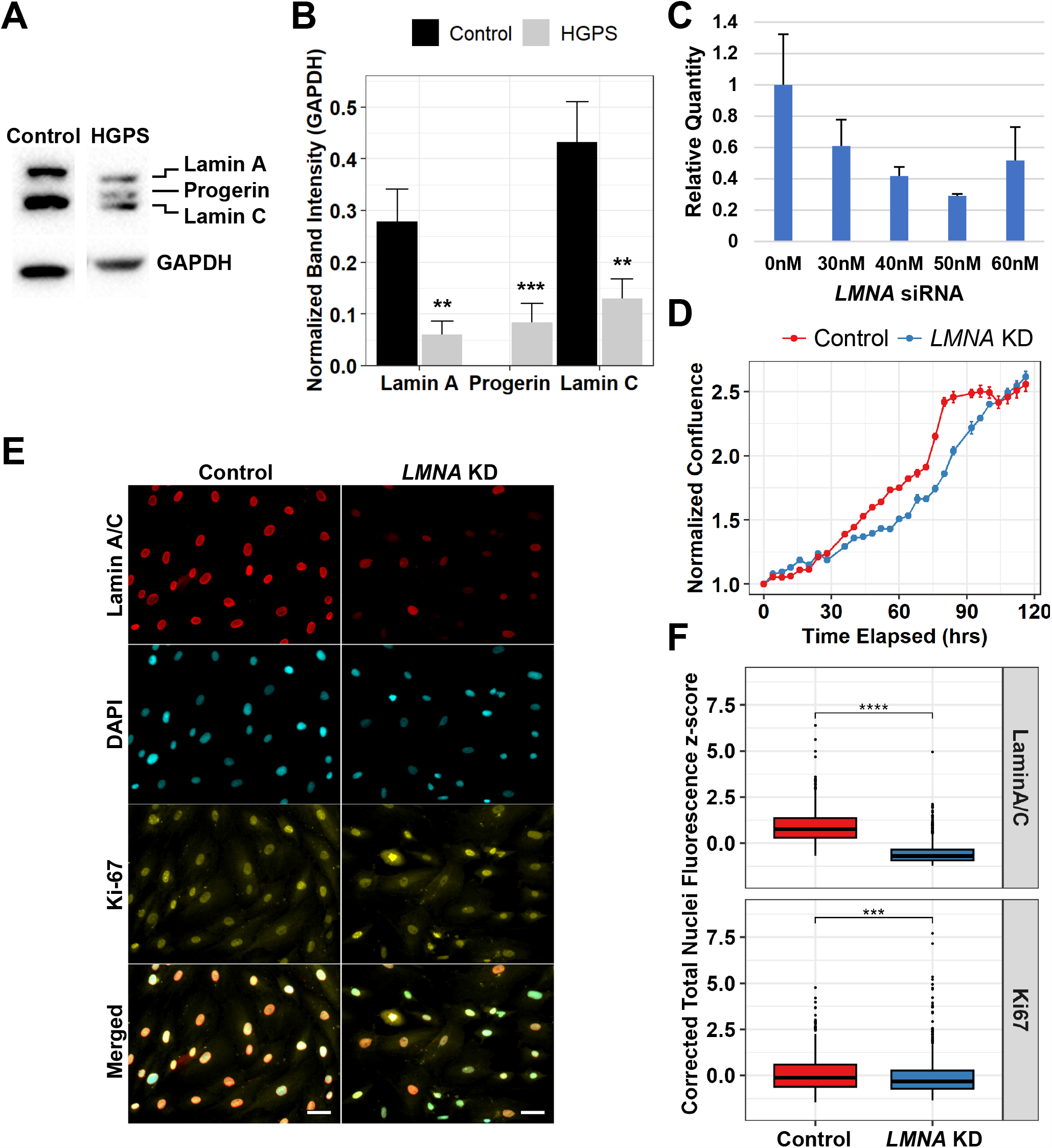
Validation of *LMNA* Knockdown. Representative western blot **A)** image and **B)** quantitation of Lamin A/C in control and HGPS fibroblast lysates normalized to GAPDH. **C)** Quantitative PCR (qPCR) assessment of *LMNA* loss 5 days after transfection with varying concentrations of *LMNA* DsiRNAs. **D)** Cell proliferation assay comparing control and *LMNA* knockdown fibroblasts over 120 hours. **E)** Representative immunofluorescence images and **F)** quantification of Ki-67 and Lamin A/C fluorescence intensity (z-scores) 5 days after *LMNA* knockdown. Scale bars are 25 μm. Statistical comparisons are unpaired Mann-Whitney U-tests. **p<0.01, ***p<0.001, ****p<0.0001.

**Supplemental Figure 2.**
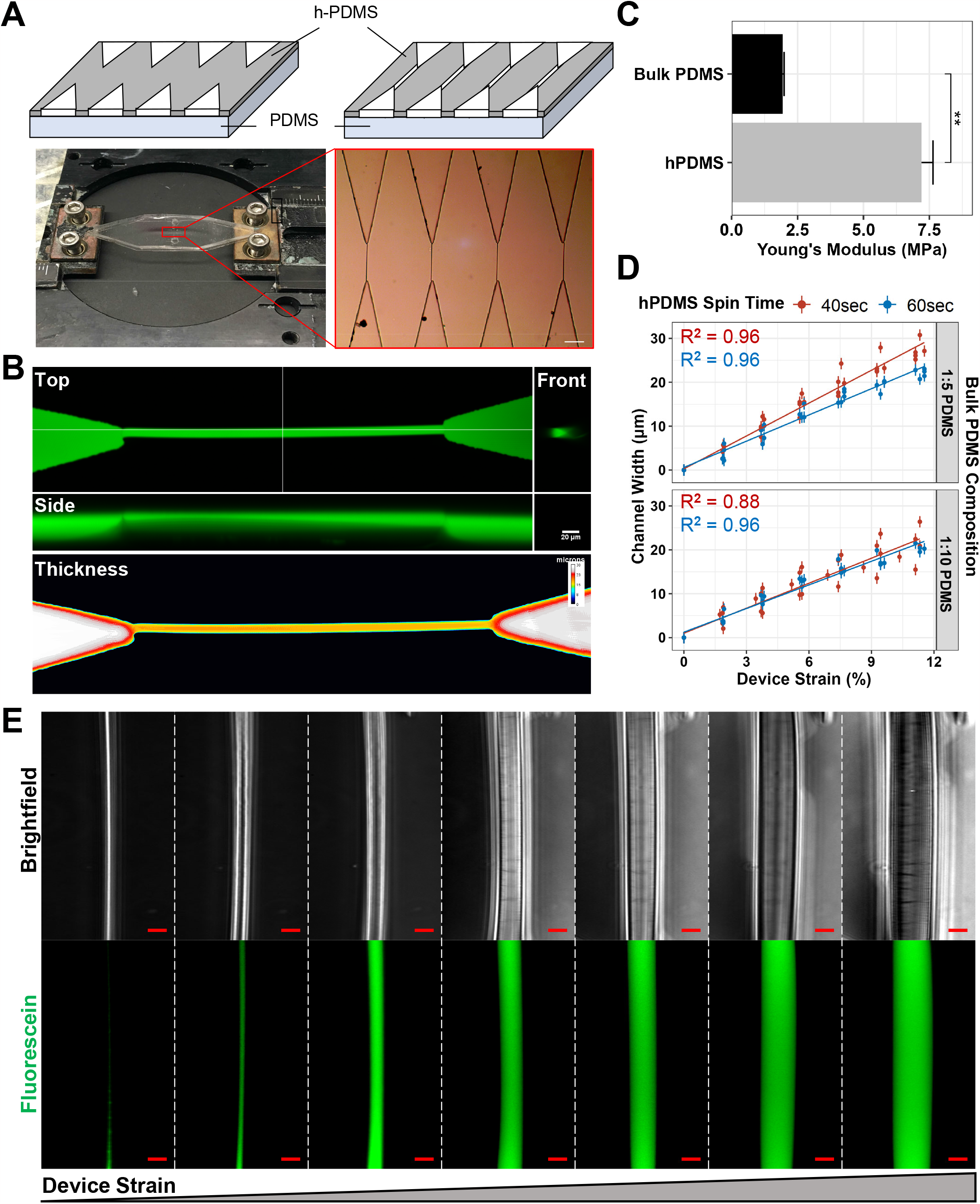
Fabrication and Characterization of Micro-Mechanical Device for Assaying Nuclear Mechanics. **A)** Schematic of device fabrication and channel formation through application of device strain. **B)** Orthogonal projections (top) and thickness heatmap (bottom) of an open fluorescein-filled microchannel. Scale bar is 20 μm. **C)** Measurement of Young’s moduli in h-PDMS and PDMS layers through strain-controlled unilateral tension. Statistical comparison is unpaired Mann-Whitney U-test. **p<0.01. **D)** Quantitation of channel widths versus device strains in devices with varying hPDMS thicknesses and bulk PDMS compositions. Data for each type of device are fitted with a regression line. **E)** Representative images of a fluorescein-filled microchannel opening under applied device strain. Scale bar is 10 μm.

**Supplemental Figure 3.**
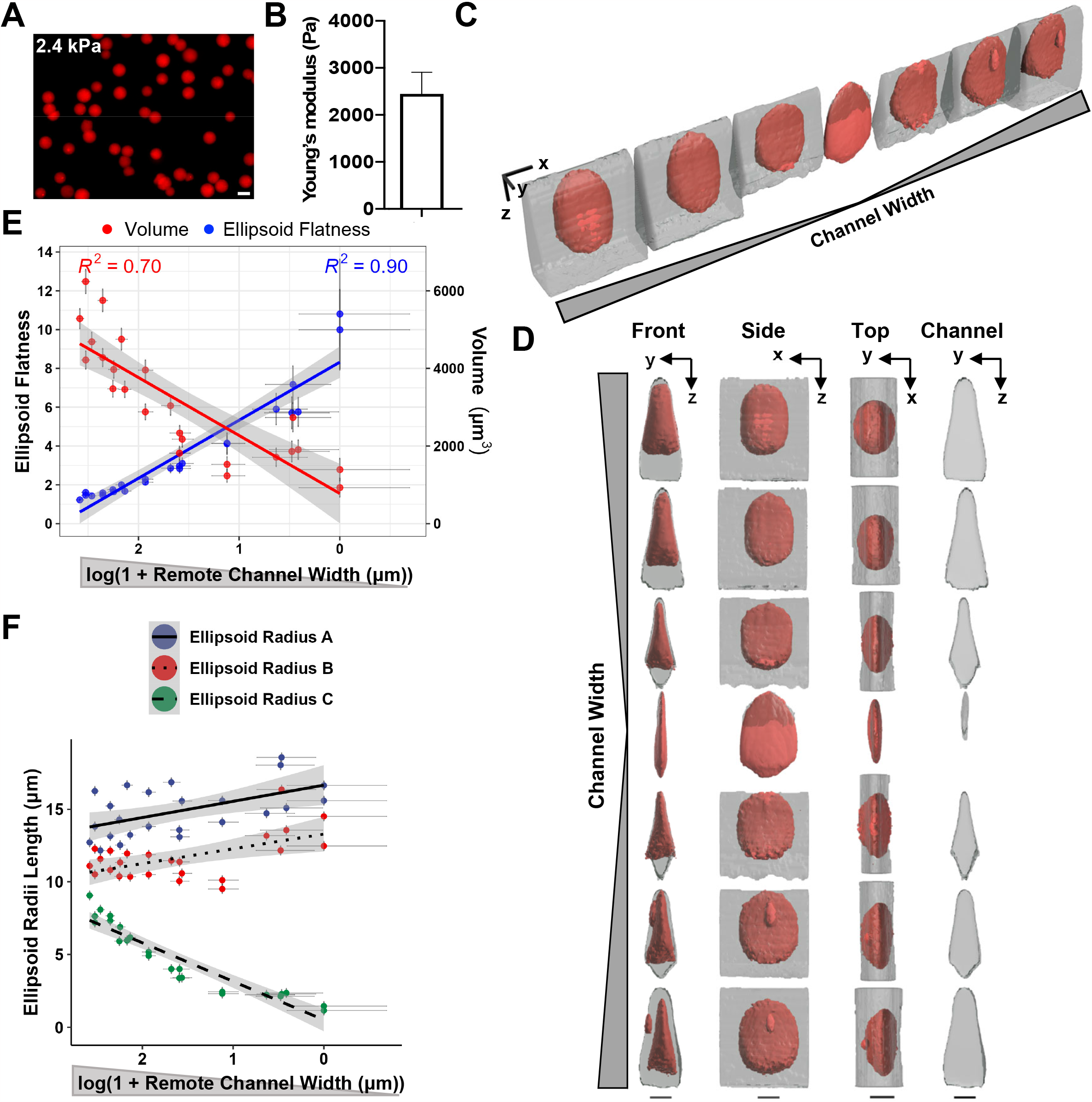
Micro-mechanical Compression of Alginate Hydrogel Beads. **A)** Representative fluorescence image of rhodamine-conjugated alginate hydrogel beads designed to have Young’s moduli equal to 2.4 kPa. Scale bar is 25 μm. **B)** Young’s modulus of hydrogel beads as measured by atomic force microscopy. Representative confocal reconstructions of **C)** angled and **D)** orthogonal views of changes in hydrogel bead (red) morphology during channel (clear) closing and opening. Scale bars (black) are 1 μm × 1 μm × 10 μm and located under the center of the beads in the orthogonal views. **E)** Flatness of fitted ellipsoids and volume of hydrogel beads during compression as a function of channel width distal to the bead. Pearson’s correlation is shown for each fit. (n=6 beads). **F)** Fitted ellipsoid radii as a function of remote channel width. Radii are ordered by decreasing length (A>B>C). (n=6 beads).

**Supplemental Figure 4.**
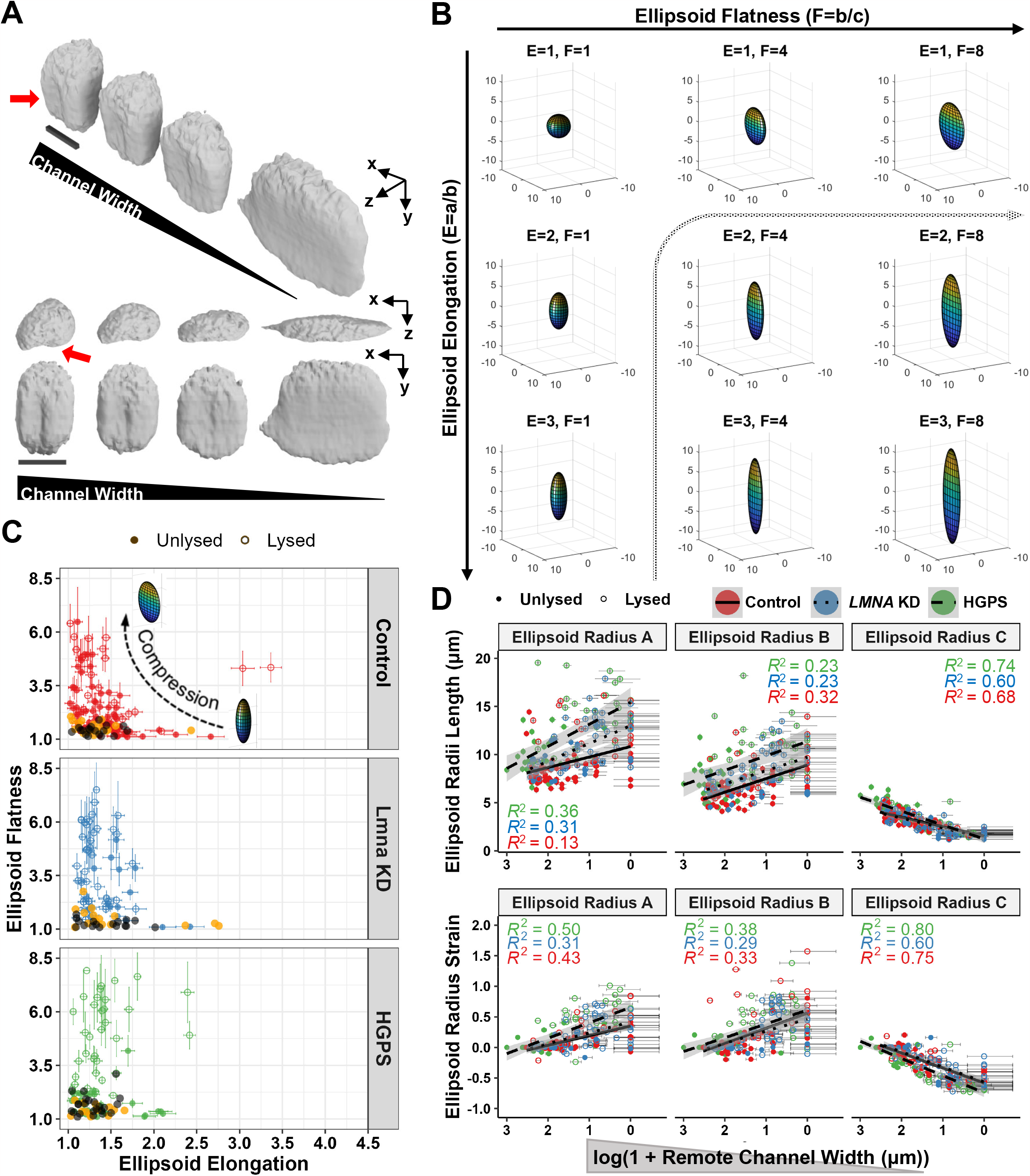
Ellipsoids Fitted to Nuclei Reveal Changes in Nuclear Morphology During Compression. **A)** Confocal reconstructions of a nuclear wrinkle being smoothed from several perspectives during compression. The wrinkle is indicated with a red arrow. Scale bars (black bars) are 1 μm × 1 μm × 10 μm. **B)** Diagram of the relationship between the ellipsoid elongation and flatness factors. The observed trend in nuclear compression data is shown as a dotted arrow through this space. **C)** Comparison of fitted ellipsoid flatness and elongation during cellular compression. Lysis status is marked by open and closed circles. Cells lying outside of the channel in the inlet and cells sitting on a glass coverslip are shown in orange and black, respectively. **D)** Radii of ellipsoids fitted to each cell type (top) and their strains (bottom) as a function of remote channel width. Ellipsoid radii are ordered by decreasing length (A>B>C). A line is fitted to data for each cell type and ellipsoid radius.

**Supplemental Figure 5.**
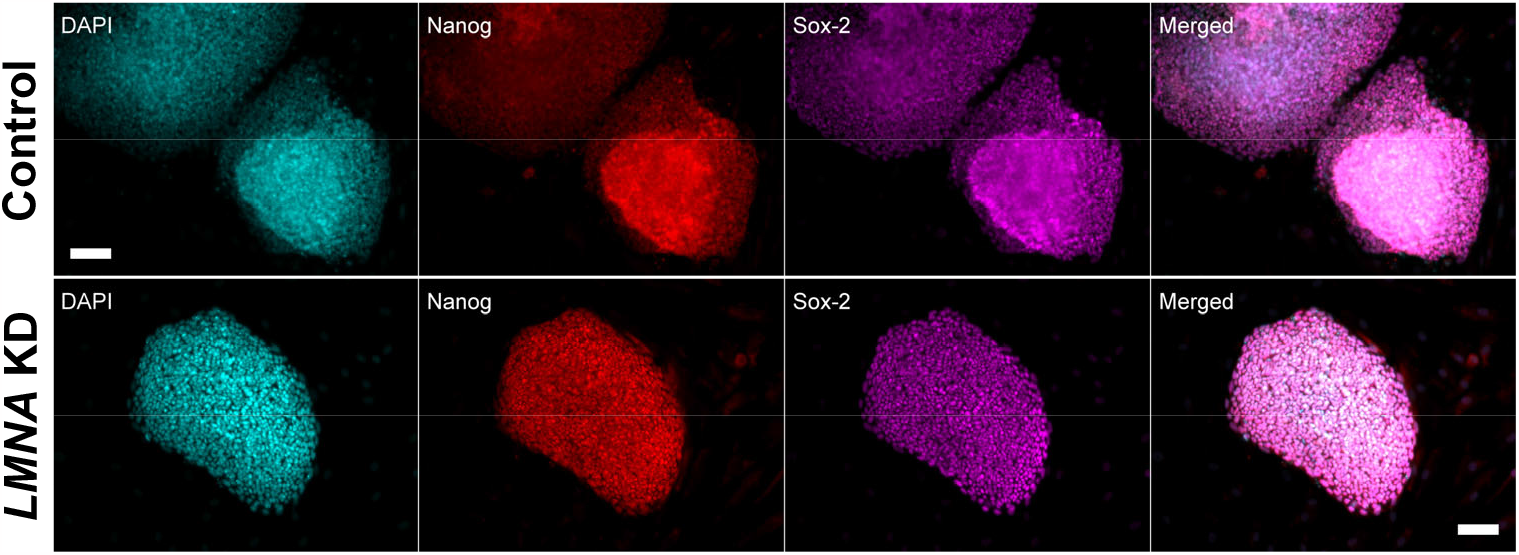
Immunofluorescence Staining for Pluripotency Markers Identifies Genuinely Reprogrammed Colonies. Representative immunofluorescence staining of Sox-2 (magenta) and Nanog (red) at days 22 after reprogramming. Scale bars are 100 μm.

**Supplemental Figure 6.**
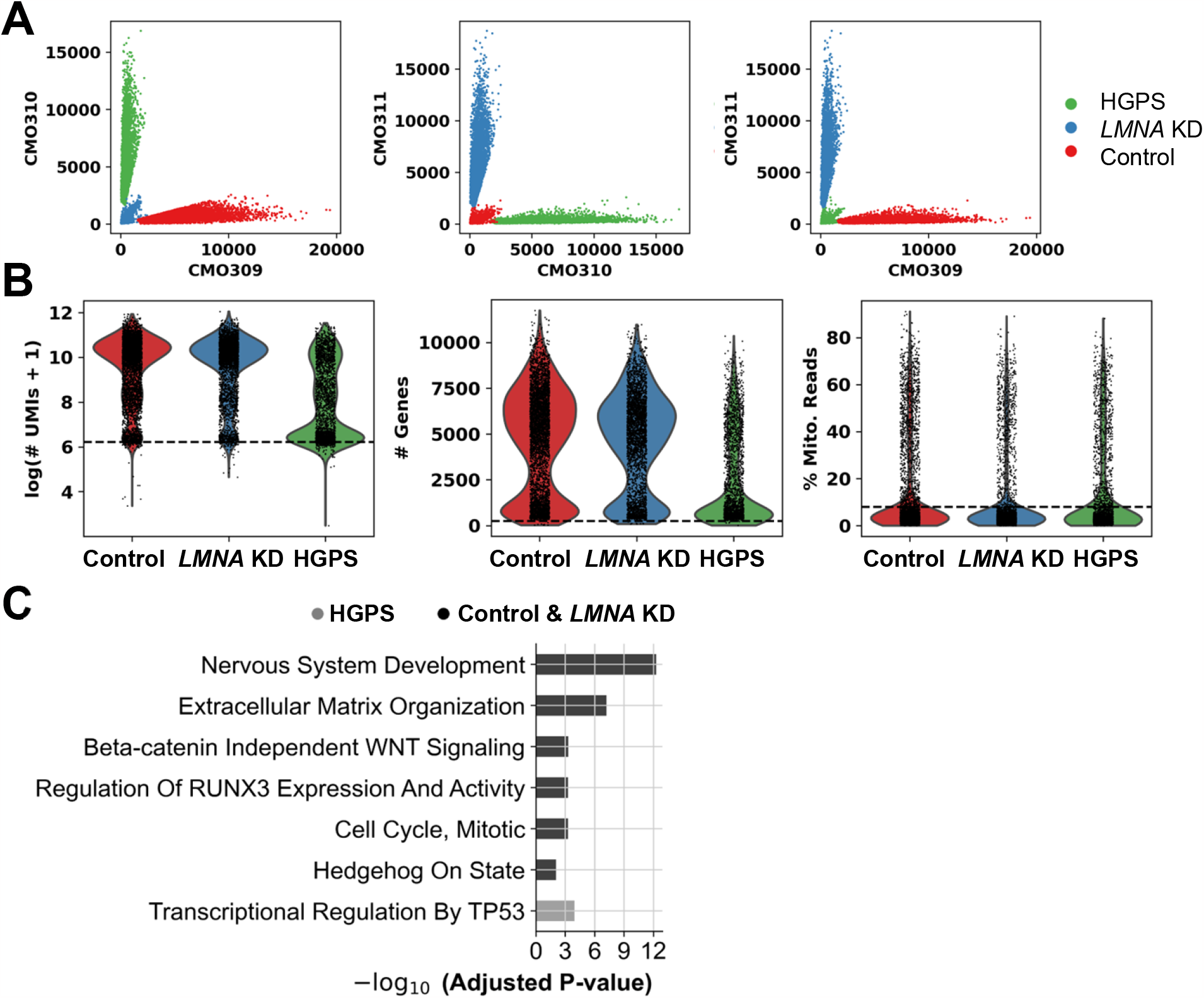
Generation and Quality Control Filtering of Single-Cell RNA-Sequencing Datasets. **A)** Confident assignments of barcodes to single cells per sample demultiplexing multiplexing oligos. **B)** Quality control metrics for combined dataset with thresholds for high-quality cells shown as dashed lines. **C)** Enriched Reactome terms from differentially expressed genes in HGPS fibroblasts compared to Normal and *LMNA* knockdown fibroblasts in Cluster 4.

## References

1. Voss, T. C. & Hager, G. L. Dynamic regulation of transcriptional states by chromatin and transcription factors. Nat Rev Genet 15, 69–81 (2014).

2. Yadav, T., Quivy, J.-P. & Almouzni, G. Chromatin plasticity: A versatile landscape that underlies cell fate and identity. Science 361, 1332–1336 (2018).

3. Misteli, T. Beyond the Sequence: Cellular Organization of Genome Function. Cell 128, 787–800 (2007).

4. van Steensel, B. & Dekker, J. Genomics tools for unraveling chromosome architecture. Nat Biotechnol 28, 1089–1095 (2010).

5. Solovei, I. et al. LBR and Lamin A/C Sequentially Tether Peripheral Heterochromatin and Inversely Regulate Differentiation. Cell 152, 584–598 (2013).

6. Falk, M. et al. Heterochromatin drives compartmentalization of inverted and conventional nuclei. Nature 570, 395–399 (2019).

7. Ho, C. Y. & Lammerding, J. Lamins at a glance. Journal of Cell Science 125, 2087–2093 (2012).

8. Adam, S. A. The Nucleoskeleton. Cold Spring Harb Perspect Biol 9, a023556 (2017).

9. Nuclear lamins are not required for lamina-associated domain organization in mouse embryonic stem cells. EMBO reports 16, 610–617 (2015).

10. Peric-Hupkes, D. et al. Molecular Maps of the Reorganization of Genome-Nuclear Lamina Interactions during Differentiation. Molecular Cell 38, 603–613 (2010).

11. Poleshko, A. et al. Genome-Nuclear Lamina Interactions Regulate Cardiac Stem Cell Lineage Restriction. Cell 171, 573-587.e14 (2017).

12. Wong, X., Luperchio, T. R. & Reddy, K. L. NET gains and losses: the role of changing nuclear envelope proteomes in genome regulation. Current Opinion in Cell Biology 28, 105–120 (2014).

13. Swift, J. et al. Nuclear Lamin-A Scales with Tissue Stiffness and Enhances Matrix-Directed Differentiation. Science 341, 1240104 (2013).

14. Robson, M. I. et al. Tissue-Specific Gene Repositioning by Muscle Nuclear Membrane Proteins Enhances Repression of Critical Developmental Genes during Myogenesis. Molecular Cell 62, 834–847 (2016).

15. van Steensel, B. & Belmont, A. S. Lamina-Associated Domains: Links with Chromosome Architecture, Heterochromatin, and Gene Repression. Cell 169, 780–791 (2017).

16. Bustin, M. & Misteli, T. Nongenetic functions of the genome. Science 352, aad6933 (2016).

17. Guelen, L. et al. Domain organization of human chromosomes revealed by mapping of nuclear lamina interactions. Nature 453, 948–951 (2008).

18. Zuleger, N., Robson, M. I. & Schirmer, E. C. The nuclear envelope as a chromatin organizer. Nucleus 2, 339–349 (2011).

19. Demmerle, J., Koch, A. J. & Holaska, J. M. Emerin and histone deacetylase 3 (HDAC3) cooperatively regulate expression and nuclear positions of MyoD, Myf5, and Pax7 genes during myogenesis. Chromosome Res 21, 765–779 (2013).

20. Robson, M. I. et al. Constrained release of lamina-associated enhancers and genes from the nuclear envelope during T-cell activation facilitates their association in chromosome compartments. Genome Res. 27, 1126–1138 (2017).

21. Merideth, M. A. et al. Phenotype and Course of Hutchinson–Gilford Progeria Syndrome. New England Journal of Medicine 358, 592–604 (2008).

22. De Sandre-Giovannoli, A. et al. Lamin A Truncation in Hutchinson-Gilford Progeria. Science 300, 2055–2055 (2003).

23. Goldman, R. D. et al. Accumulation of mutant lamin A causes progressive changes in nuclear architecture in Hutchinson–Gilford progeria syndrome. PNAS 101, 8963–8968 (2004).

24. Isermann, P. & Lammerding, J. Nuclear Mechanics and Mechanotransduction in Health and Disease. Current Biology 23, R1113–R1121 (2013).

25. Swift, J. & Discher, D. E. The nuclear lamina is mechano-responsive to ECM elasticity in mature tissue. Journal of Cell Science 127, 3005–3015 (2014).

26. Lammerding, J. et al. Lamins A and C but Not Lamin B1 Regulate Nuclear Mechanics. Journal of Biological Chemistry 281, 25768–25780 (2006).

27. Stephens, A. D., Banigan, E. J., Adam, S. A., Goldman, R. D. & Marko, J. F. Chromatin and lamin A determine two different mechanical response regimes of the cell nucleus. MBoC 28, 1984–1996 (2017).

28. Harada, T. et al. Nuclear lamin stiffness is a barrier to 3D migration, but softness can limit survival. J Cell Biol 204, 669–682 (2014).

29. Cacchiarelli, D. et al. Integrative analyses of human reprogramming reveal dynamic nature of induced pluripotency. Cell 162, 412–424 (2015).

30. Scaffidi, P. & Misteli, T. Reversal of the cellular phenotype in the premature aging disease Hutchinson-Gilford progeria syndrome. Nat Med 11, 440–445 (2005).

31. Kim, D.-H. et al. Synthetic dsRNA Dicer substrates enhance RNAi potency and efficacy. Nat Biotechnol 23, 222–226 (2005).

32. Poleshko, A. et al. The Human Protein PRR14 Tethers Heterochromatin to the Nuclear Lamina during Interphase and Mitotic Exit. Cell Reports 5, 292–301 (2013).

33. Beagan, J. A. et al. Local Genome Topology Can Exhibit an Incompletely Rewired 3D-Folding State during Somatic Cell Reprogramming. Cell Stem Cell 18, 611–624 (2016).

34. Köhler, F. et al. Epigenetic deregulation of lamina-associated domains in Hutchinson-Gilford progeria syndrome. Genome Medicine 12, 46 (2020).

35. Buenrostro, J. D., Giresi, P. G., Zaba, L. C., Chang, H. Y. & Greenleaf, W. J. Transposition of native chromatin for fast and sensitive epigenomic profiling of open chromatin, DNA-binding proteins and nucleosome position. Nature Methods 10, 1213–1218 (2013).

36. Corces, M. R. et al. An improved ATAC-seq protocol reduces background and enables interrogation of frozen tissues. Nat Methods 14, 959–962 (2017).

37. McLean, C. Y. et al. GREAT improves functional interpretation of cis-regulatory regions. Nature Biotechnology 28, 495–501 (2010).

38. Kraushaar, D. C., Dalton, S. & Wang, L. Heparan sulfate: a key regulator of embryonic stem cell fate. Biological Chemistry 394, 741–751 (2013).

39. Cappuccio, I. et al. Endogenous activation of mGlu5 metabotropic glutamate receptors supports selfrenewal of cultured mouse embryonic stem cells. Neuropharmacology 49, 196–205 (2005).

40. Spinsanti, P. et al. Endogenously activated mGlu5 metabotropic glutamate receptors sustain the increase in c-Myc expression induced by leukaemia inhibitory factor in cultured mouse embryonic stem cells. Journal of Neurochemistry 99, 299–307 (2006).

41. Pauklin, S. & Vallier, L. Activin/Nodal signalling in stem cells. Development 142, 607–619 (2015).

42. Zhu, Y., Fan, C. & Zhao, B. Differential expression of piRNAs in reprogrammed pluripotent stem cells from mouse embryonic fibroblasts. IUBMB Life 71, 1906–1915 (2019).

43. Pezic, D., Manakov, S. A., Sachidanandam, R. & Aravin, A. A. piRNA pathway targets active LINE1 elements to establish the repressive H3K9me3 mark in germ cells. Genes Dev. 28, 1410–1428 (2014).

44. Bentsen, M. et al. ATAC-seq footprinting unravels kinetics of transcription factor binding during zygotic genome activation. Nat Commun 11, 4267 (2020).

45. Barrallo-Gimeno, A. & Nieto, M. A. The Snail genes as inducers of cell movement and survival: implications in development and cancer. Development 132, 3151–3161 (2005).

46. Tahmasebi, S. et al. Control of embryonic stem cell self-renewal and differentiation via coordinated alternative splicing and translation of YY2. Proceedings of the National Academy of Sciences 113, 12360–12367 (2016).

47. Nirala, N. K. et al. Hinfp is a guardian of the somatic genome by repressing transposable elements. Proceedings of the National Academy of Sciences 118, e2100839118 (2021).

48. Markov, G. J. et al. AP-1 is a temporally regulated dual gatekeeper of reprogramming to pluripotency. Proceedings of the National Academy of Sciences 118, e2104841118 (2021).

49. Chronis, C. et al. Cooperative Binding of Transcription Factors Orchestrates Reprogramming. Cell 168, 442-459.e20 (2017).

50. Xing, Q. R. et al. Diversification of reprogramming trajectories revealed by parallel single-cell transcriptome and chromatin accessibility sequencing. Science Advances 6, eaba1190 (2020).

51. Huang, X. et al. A chromodomain protein mediates heterochromatin-directed piRNA expression. Proceedings of the National Academy of Sciences 118, e2103723118 (2021).

52. Iwasaki, Y. W. et al. Piwi–piRNA complexes induce stepwise changes in nuclear architecture at target loci. The EMBO Journal 40, e108345 (2021).

53. Meuleman, W. et al. Constitutive nuclear lamina–genome interactions are highly conserved and associated with A/T-rich sequence. Genome Res. 23, 270–280 (2013).

54. McGarry, J. P. Characterization of Cell Mechanical Properties by Computational Modeling of Parallel Plate Compression. Ann Biomed Eng 37, 2317–2325 (2009).

55. Massou, S. et al. Cell stretching is amplified by active actin remodelling to deform and recruit proteins in mechanosensitive structures. Nat Cell Biol 22, 1011–1023 (2020).

56. Nava, M. M. et al. Heterochromatin-Driven Nuclear Softening Protects the Genome against Mechanical Stress-Induced Damage. Cell 181, 800-817.e22 (2020).

57. Hobson, C. M. et al. Correlating nuclear morphology and external force with combined atomic force microscopy and light sheet imaging separates roles of chromatin and lamin A/C in nuclear mechanics. MBoC 31, 1788–1801 (2020).

58. Liu, H. et al. In Situ Mechanical Characterization of the Cell Nucleus by Atomic Force Microscopy. ACS Nano 8, 3821–3828 (2014).

59. Lomakin, A. J. et al. The nucleus acts as a ruler tailoring cell responses to spatial constraints. Science 370, eaba2894 (2020).

60. Dahl, K. N. et al. Distinct structural and mechanical properties of the nuclear lamina in Hutchinson-Gilford progeria syndrome. Proceedings of the National Academy of Sciences 103, 10271–10276 (2006).

61. Pushkarsky, I. et al. Elastomeric sensor surfaces for high-throughput single-cell force cytometry. Nat Biomed Eng 2, 124–137 (2018).

62. Ho, K. K. Y., Wang, Y. L., Wu, J. & Liu, A. P. Advanced Microfluidic Device Designed for Cyclic Compression of Single Adherent Cells. Front. Bioeng. Biotechnol. 6, (2018).

63. Zhu, X. et al. Fabrication of reconfigurable protein matrices by cracking. Nature Materials 4, 403–406 (2005).

64. Kim, B. C. et al. Fracture-Based Fabrication of Normally Closed, Adjustable, and Fully Reversible Microscale Fluidic Channels. Small 10, 4020–4029 (2014).

65. Mao, A. S. et al. Deterministic encapsulation of single cells in thin tunable microgels for niche modelling and therapeutic delivery. Nature Materials 16, 236–243 (2017).

66. Kalukula, Y., Stephens, A. D., Lammerding, J. & Gabriele, S. Mechanics and functional consequences of nuclear deformations. Nat Rev Mol Cell Biol 23, 583–602 (2022).

67. Takahashi, K. & Yamanaka, S. Induction of Pluripotent Stem Cells from Mouse Embryonic and Adult Fibroblast Cultures by Defined Factors. Cell 126, 663–676 (2006).

68. Tanabe, K., Nakamura, M., Narita, M., Takahashi, K. & Yamanaka, S. Maturation, not initiation, is the major roadblock during reprogramming toward pluripotency from human fibroblasts. Proceedings of the National Academy of Sciences 110, 12172–12179 (2013).

69. Chen, Z. et al. Reprogramming progeria fibroblasts re-establishes a normal epigenetic landscape. Aging Cell 16, 870–887 (2017).

70. McInnes, L., Healy, J. & Melville, J. UMAP: Uniform Manifold Approximation and Projection for Dimension Reduction. arXiv:1802.03426 [cs, stat] (2018).

71. Wolf, F. A. et al. PAGA: graph abstraction reconciles clustering with trajectory inference through a topology preserving map of single cells. Genome Biology 20, 59 (2019).

72. Haghverdi, L., Büttner, M., Wolf, F. A., Buettner, F. & Theis, F. J. Diffusion pseudotime robustly reconstructs lineage branching. Nat Methods 13, 845–848 (2016).

73. Jiao, J. et al. Promoting Reprogramming by FGF2 Reveals that the Extracellular Matrix Is a Barrier for Reprogramming Fibroblasts to Pluripotency. Stem Cells 31, 729–740 (2013).

74. Miki, T., Yasuda, S. & Kahn, M. Wnt/β-catenin Signaling in Embryonic Stem Cell Self-renewal and Somatic Cell Reprogramming. Stem Cell Rev and Rep 7, 836–846 (2011).

75. Wu, S. M., Choo, A. B. H., Yap, M. G. S. & Chan, K. K.-K. Role of Sonic hedgehog signaling and the expression of its components in human embryonic stem cells. Stem Cell Research 4, 38–49 (2010).

76. Dannenmann, B. et al. High Glutathione and Glutathione Peroxidase-2 Levels Mediate Cell-Type-Specific DNA Damage Protection in Human Induced Pluripotent Stem Cells. Stem Cell Reports 4, 886–898 (2015).

77. Iratni, R. et al. Inhibition of Excess Nodal Signaling During Mouse Gastrulation by the Transcriptional Corepressor DRAP1. Science 298, 1996–1999 (2002).

78. Mor, N. et al. Neutralizing Gatad2a-Chd4-Mbd3/NuRD Complex Facilitates Deterministic Induction of Naive Pluripotency. Cell Stem Cell 23, 412-425.e10 (2018).

79. Botti, V. et al. Cellular differentiation state modulates the mRNA export activity of SR proteins. Journal of Cell Biology 216, 1993–2009 (2017).

80. Pagliara, S. et al. Auxetic nuclei in embryonic stem cells exiting pluripotency. Nature Mater 13, 638–644 (2014).

81. Roy, B. et al. Laterally confined growth of cells induces nuclear reprogramming in the absence of exogenous biochemical factors. Proceedings of the National Academy of Sciences 115, E4741–E4750 (2018).

82. Song, Y. et al. Transient nuclear deformation primes epigenetic state and promotes cell reprogramming. Nat. Mater. 1–9 (2022) doi:10.1038/s41563-022-01312-3.

83. Broers, J. L. V. et al. Decreased mechanical stiffness in LMNA−/− cells is caused by defective nucleo-cytoskeletal integrity: implications for the development of laminopathies. Human Molecular Genetics 13, 2567–2580 (2004).

84. Poleshko, A. et al. H3K9me2 orchestrates inheritance of spatial positioning of peripheral heterochromatin through mitosis. eLife 8, e49278 (2019).

85. Nicetto, D. et al. H3K9me3-heterochromatin loss at protein-coding genes enables developmental lineage specification. Science 363, 294–297 (2019).

86. Methot, S. P. et al. H3K9me selectively blocks transcription factor activity and ensures differentiated tissue integrity. Nat Cell Biol 23, 1163–1175 (2021).

87. Esteban, M. A. et al. Vitamin C Enhances the Generation of Mouse and Human Induced Pluripotent Stem Cells. Cell Stem Cell 6, 71–79 (2010).

88. Ebata, K. T. et al. Vitamin C induces specific demethylation of H3K9me2 in mouse embryonic stem cells via Kdm3a/b. Epigenetics & Chromatin 10, 36 (2017).

89. Shumaker, D. K. et al. Mutant nuclear lamin A leads to progressive alterations of epigenetic control in premature aging. Proceedings of the National Academy of Sciences 103, 8703–8708 (2006).

90. McCord, R. P. et al. Correlated alterations in genome organization, histone methylation, and DNA– lamin A/C interactions in Hutchinson-Gilford progeria syndrome. Genome Res. 23, 260–269 (2013).

91. Ikegami, K., Secchia, S., Almakki, O., Lieb, J. D. & Moskowitz, I. P. Phosphorylated Lamin A/C in the Nuclear Interior Binds Active Enhancers Associated with Abnormal Transcription in Progeria. Developmental Cell 52, 699-713.e11 (2020).

92. Sebestyén, E. et al. SAMMY-seq reveals early alteration of heterochromatin and deregulation of bivalent genes in Hutchinson-Gilford Progeria Syndrome. Nat Commun 11, 6274 (2020).

93. Chandra, T. et al. Global Reorganization of the Nuclear Landscape in Senescent Cells. Cell Reports 10, 471–483 (2015).

94. Lammerding, J., Engler, A. J. & Kamm, R. Mechanobiology of the cell nucleus. APL Bioengineering 6, 040401 (2022).

95. Fan, Y.-J. et al. Microfluidic channel integrated with a lattice lightsheet microscopic system for continuous cell imaging. Lab Chip 21, 344–354 (2021).

96. Cosgrove, B. D. et al. Nuclear envelope wrinkling predicts mesenchymal progenitor cell mechano-response in 2D and 3D microenvironments. Biomaterials 270, 120662 (2021).

97. Hasper, J. et al. Long lifetime and selective accumulation of the A-type lamins accounts for the tissue specificity of Hutchinson-Gilford progeria syndrome. 2023.02.04.527139 Preprint at https://doi.org/10.1101/2023.02.04.527139 (2023).

98. Pajerowski, J. D., Dahl, K. N., Zhong, F. L., Sammak, P. J. & Discher, D. E. Physical plasticity of the nucleus in stem cell differentiation. PNAS 104, 15619–15624 (2007).

99. Basu, A. & Tiwari, V. K. Epigenetic reprogramming of cell identity: lessons from development for regenerative medicine. Clinical Epigenetics 13, 144 (2021).

100. Lenain, C. et al. Massive reshaping of genome–nuclear lamina interactions during oncogene-induced senescence. Genome Res. 27, 1634–1644 (2017).

101. Maher, B. ENCODE: The human encyclopaedia. Nature 489, 46–48 (2012).

102. Yu, G., Wang, L.-G. & He, Q.-Y. ChIPseeker: an R/Bioconductor package for ChIP peak annotation, comparison and visualization. Bioinformatics 31, 2382–2383 (2015).

103. McGinnis, C. S., Murrow, L. M. & Gartner, Z. J. DoubletFinder: Doublet Detection in Single-Cell RNA Sequencing Data Using Artificial Nearest Neighbors. Cell Systems 8, 329-337.e4 (2019).

104. Ramírez, F. et al. deepTools2: a next generation web server for deep-sequencing data analysis. Nucleic Acids Research 44, W160–W165 (2016).

105. Castro-Mondragon, J. A. et al. JASPAR 2022: the 9th release of the open-access database of transcription factor binding profiles. Nucleic Acids Research 50, D165–D173 (2022).

106. Liao, Y., Wang, J., Jaehnig, E. J., Shi, Z. & Zhang, B. WebGestalt 2019: gene set analysis toolkit with revamped UIs and APIs. Nucleic Acids Research 47, W199–W205 (2019).

107. Wolf, F. A., Angerer, P. & Theis, F. J. SCANPY: large-scale single-cell gene expression data analysis. Genome Biology 19, 15 (2018).

108. Fang, Z., Liu, X. & Peltz, G. GSEApy: a comprehensive package for performing gene set enrichment analysis in Python. Bioinformatics 39, btac757 (2023).

109. Legland, D., Arganda-Carreras, I. & Andrey, P. MorphoLibJ: integrated library and plugins for mathematical morphology with ImageJ. Bioinformatics 32, 3532–3534 (2016).

110. Bolte, S. & Cordelières, F. P. A guided tour into subcellular colocalization analysis in light microscopy. Journal of Microscopy 224, 213–232 (2006).

111. Schmid, B., Schindelin, J., Cardona, A., Longair, M. & Heisenberg, M. A high-level 3D visualization API for Java and ImageJ. BMC Bioinformatics 11, 274 (2010).

112. Ollion, J., Cochennec, J., Loll, F., Escudé, C. & Boudier, T. TANGO: a generic tool for highthroughput 3D image analysis for studying nuclear organization. Bioinformatics 29, 1840–1841 (2013).

113. Yang, B. A. et al. Sestrins regulate muscle stem cell metabolic homeostasis. Stem Cell Reports 0, (2021).

114. Kim, B. C. et al. Guided fracture of films on soft substrates to create micro/nano-feature arrays with controlled periodicity. Sci Rep 3, 3027 (2013).

115. Yang, B. A., Westerhof, T. M., Sabin, K., Merajver, S. D. & Aguilar, C. A. Engineered Tools to Study Intercellular Communication. Advanced Science 8, 2002825 (2021).

116. Schmid, H. & Michel, B. Siloxane Polymers for High-Resolution, High-Accuracy Soft Lithography. Macromolecules 33, 3042–3049 (2000).

117. Choul Kim, B., Moraes, C., Huang, J., D. Thouless, M. & Takayama, S. Fracture-based micro- and nanofabrication for biological applications. Biomaterials Science 2, 288–296 (2014).

118. Ritchie, M. E. et al. limma powers differential expression analyses for RNA-sequencing and microarray studies. Nucleic Acids Res 43, e47–e47 (2015).

119. Shcherbina, A. et al. Dissecting Murine Muscle Stem Cell Aging through Regeneration Using Integrative Genomic Analysis. Cell Reports 32, 107964 (2020).

120. Amemiya, H. M., Kundaje, A. & Boyle, A. P. The ENCODE Blacklist: Identification of Problematic Regions of the Genome. Scientific Reports 9, 9354 (2019).

121. Kuleshov, M. V. et al. Enrichr: a comprehensive gene set enrichment analysis web server 2016 update. Nucleic Acids Research 44, W90–W97 (2016).

122. Jacomy, M., Venturini, T., Heymann, S. & Bastian, M. ForceAtlas2, a Continuous Graph Layout Algorithm for Handy Network Visualization Designed for the Gephi Software. PLOS ONE 9, e98679 (2014).

